# EPRS Regulates Proline-rich Pro-fibrotic Protein Synthesis during Cardiac Fibrosis

**DOI:** 10.1101/777490

**Authors:** Jiangbin Wu, Kadiam C Venkata Subbaiah, Li Huitong Xie, Feng Jiang, Deanne Mickelsen, Jason R Myers, Wai Hong Wilson Tang, Peng Yao

**Author notes:** Correspondence to Peng Yao, Ph.D., Aab Cardiovascular Research Institute, Department of Medicine, University of Rochester School of Medicine & Dentistry, Rochester, New York 14586. Abbreviations of key terms:* ACM, adult cardiomyocytes; ARS, aminoacyl-tRNA synthetase; CF, cardiac fibroblasts; CM, cardiomyocytes; CVD, cardiovascular disease; ECM, extracellular matrix; EPRS, glutamyl-prolyl-tRNA synthetase; Halo, halofuginone; HF, heart failure; ISO, isoproterenol; Pro-tRNA^Pro^, prolyl tRNA^Pro^, proline ligated to tRNA^Pro^; PRR, proline-rich (XPPY codon motifs); RBP, RNA-binding protein; TAC, transverse aortic constriction; TMX, tamoxifen.

## Abstract

**Rationale:** Increased protein synthesis of pro-fibrotic genes is a common feature of cardiac fibrosis, a major manifestation of heart failure. Despite this important observation, critical factors and molecular mechanisms for translational control of pro-fibrotic genes during cardiac fibrosis remain unclear.

**Objective:** This study aimed to test the hypothesis that cardiac stress-induced expression of a bifunctional aminoacyl-tRNA synthetase (ARS), glutamyl-prolyl-tRNA synthetase (EPRS), is preferentially required for the translation of proline codon-rich (PRR) pro-fibrotic mRNAs in cardiac fibroblasts during cardiac fibrosis.

**Methods and Results:** By analyses of multiple available unbiased large-scale screening datasets of human and mouse heart failure, we have discovered that EPRS acts as an integrated node among all the ARSs in various cardiac pathogenic processes. We confirmed that EPRS was induced at both mRNA and protein level (∼1.5-2.5 fold increase) in failing hearts compared with non-failing hearts using our cohort of human and mouse heart samples. Genetic knockout of one allele of *Eprs* globally (*Eprs*^+/-^) using CRISPR-Cas9 technology or in a myofibroblast-specific manner (*Eprs*^flox/+^; *Postn*^MCM/+^) strongly reduces cardiac fibrosis (∼50% reduction) in isoproterenol- and transverse aortic constriction-induced heart failure mouse models. Inhibition of EPRS by a prolyl-tRNA synthetase (PRS)-specific inhibitor, halofuginone (Halo), significantly decreased the translation efficiency of proline-rich collagens in cardiac fibroblasts. Furthermore, using transcriptome-wide RNA-Seq and polysome profiling-Seq in Halo-treated fibroblasts, we identified multiple novel Pro-rich genes in addition to collagens, such as Ltbp2 and Sulf1, which are translationally regulated by EPRS. As a major EPRS downstream effector, SULF1 is highly enriched in human and mouse myofibroblast. siRNA-mediated knockdown of SULF1 attenuates cardiac myofibroblast activation and collagen deposition.

**Conclusions:** Our results indicate that EPRS preferentially controls the translational activation of proline codon-rich pro-fibrotic genes in cardiac fibroblasts and augments pathological cardiac remodeling.

**Novelty and Significance:** *What is known?:*

- TGF-β and IL-11 increase synthesis of pro-fibrotic proteins during cardiac fibrosis.
- Many pro-fibrotic genes contain Pro genetic codon rich motifs such as collagens.
- EPRS is an essential house-keeping enzyme required for ligating Pro to tRNA^Pro^ for the synthesis of Pro-containing proteins.

*What New Information Does This Article Contribute?:*

- This study is a pioneering investigation of translational control mechanisms of pro-fibrotic gene expression in cardiac fibrosis.
- EPRS mRNA and protein expression are induced in failing human hearts and mouse hearts undergoing pathological cardiac remodeling.
- The first demonstration of the in vivo function of EPRS in cardiac remodeling. Heterozygous Eprs global knockout and myofibroblast-specific tamoxifen-inducible Eprs conditional knockout mice show reduced pathological cardiac fibrosis under stress, suggesting that the reduction of EPRS is cardioprotective.
- Identification of novel preferential translational target genes of EPRS. We found that EPRS regulates translation of Pro-rich (PRR) transcripts, which comprise most of the ECM and secretory signaling molecules. Among those targets, we identified multiple novel PRR genes such as LTBP2 and SULF1.
- SULF1 is validated as a myofibroblast marker protein in human and mouse heart failure and a potential anti-fibrosis target gene. In cardiac fibroblasts, the synthesis of pro-fibrotic proteins is upregulated by cardiac stressors to activate extracellular matrix deposition and impair cardiac function. In this study, we have discovered an EPRS-PRR gene axis that influences translational homeostasis of pro-fibrotic proteins and promotes pathological cardiac remodeling and fibrosis. EPRS is identified as a common node downstream of multiple cardiac stressors and a novel regulatory factor that facilitates pro-fibrotic mRNA translation in cardiac fibrosis. Global and myofibroblast-specific genetic ablation of EPRS can effectively reduce cardiac fibrosis. This study reveals a novel translational control mechanism that modulates cardiac fibrosis and heart function. Mild inhibition of PRR mRNA translation could be a general therapeutic strategy for the treatment of heart disease. These findings provide novel insights into the translational control mechanisms of cardiac fibrosis and will promote the development of novel therapeutics by inhibiting pro-fibrotic translation factors or their downstream effectors.

## Introduction

Cardiovascular disease (CVD) is the leading cause of morbidity and mortality worldwide. Heart failure (HF), a major manifestation of CVD, is often accompanied by cardiac fibrosis driven by increased pro-fibrotic protein synthesis following cardiac fibroblast (CF) activation^1^. According to the central dogma, protein synthesis requires gene transcription and mRNA translation, both of which can be regulated to control protein production. Transcriptional regulation has been extensively studied during cardiac fibrosis^1, 2^. However, the fact that protein expression often does not correlate with mRNA abundances^3, 4^ indicates translational control as another critical layer of regulation in promoting pro-fibrotic protein synthesis^5, 6^. Although increased pro-fibrotic protein synthesis has been observed in cardiac fibrosis^4, 5^, the underlying regulatory mechanism of pro-fibrotic mRNA translation in CFs has not been identified. This unaddressed question represents a critical gap in our understanding of the regulatory control of cardiac fibrosis in the pathological remodeling process.

The human translation machinery is comprised of three major parts, ribosomes, translation factors (initiation/elongation/termination factors), and aminoacyl-transfer RNAs (aa-tRNAs)^7^, ligation products of amino acids and tRNAs that are catalyzed by aminoacyl-tRNA synthetases (ARSs)^8, 9^. Among all mammalian ARSs, glutamyl-prolyl-tRNA synthetase (EPRS) catalyzes the attachment of two amino acids, glutamic acid (E) and proline (P), to their cognate tRNAs for protein synthesis through two catalytic domains^10^. Since plenty of pro-fibrotic proteins are proline-rich such as collagens, EPRS-mediated translation regulation probably plays a critical role in pro-fibrotic protein synthesis during cardiac fibrosis. As supporting evidence in human genetics, hypoactive mutations in the PRS domain of EPRS lead to hypomyelinating leukodystrophy without causing any known cardiac dysfunction in patients^11^. These EPRS mutations imply that reduced PRS enzymatic activity may not have adverse effects on normal heart function while reducing proline-rich protein synthesis. However, the role of EPRS in cardiac disease is still unexplored. Besides, an (E)PRS-specific inhibitor, halofuginone (Halo), blocks binding of (E)PRS to proline and tRNA^Pro^ and prevents their ligation^12–14^. Halo has been used to treat Duchenne Muscular Dystrophy (DMD) (Akashi Therapeutics) by reducing fibrosis and increasing muscle strength in phase II clinical trials. Also, GSK company confirmed its anti-fibrotic and cardiac protective activity in multiple mouse heart failure models^15^. However, the therapeutic mechanism of Halo has only been studied at the transcriptional level in triggering an amino acid starvation response (AAR) and a noncanonical TGF-β signaling pathway^12, 15, 16^. The direct downstream translational targets of EPRS by Halo inhibition remains unclear. A previous study has revealed that decreased level of ribosome, a general translation machinery component, selectively regulated translation of a subset of transcripts in human hematopoiesis^17^. Therefore, as a general translation factor, EPRS may also have its preferential translational targets during cardiac fibrosis, which has not been explored^18, 19^.

Here we examined EPRS-mediated regulatory mechanisms of pro-fibrotic protein synthesis at the translatome-wide level in fibroblasts. We show that EPRS is induced in human and mouse failing hearts. EPRS is an integrated node downstream of various cardiac pathological cues for translational control in cardiac fibrosis. Increased EPRS expression contributes to elevated translation of proline (Pro)-rich (PRR) mRNAs via enhanced translation elongation in CFs. Genetic knockout of *Eprs* antagonized cardiac pathological remodeling and fibrosis in different mouse heart failure models. Moreover, by using the EPRS inhibitor Halo, we found that EPRS inhibition selectively reduces PRR mRNA translation, such as collagens and other novel PRR genes, including LTBP2 and SULF1. Finally, we show that SULF1 is a novel biomarker for cardiac fibrosis and knockdown of SULF1 remarkably abolishes myofibroblast activation.

## Methods

### Human Specimens

All human samples of frozen cardiac tissues, including 17 samples from explanted failing hearts and 8 samples from non-failing donor hearts, as well as paraffin-embedded section slides from dilated cardiomyopathy (DCM), ischemic heart failure (IHF) or non-failing donor (NF) hearts, were acquired from the Cleveland Clinic. This study was approved by the Material Transfer Agreement between the URMC and the Cleveland Clinic.

### Mice

The *Eprs*^+/–^ global knockout chimera founder mice were produced in the University of Rochester Mouse Genome Editing Resource. We generated the *Eprs* targeted male chimera mouse in the C57BL/6J background and performed germline transmission. Using CRISPR-Cas9 system, a gRNA (GCUAGAAUUGCAACUACGUCUGG) and a homology-directed recombination DNA template containing an insertion of tandem stop codons plus an adenosine residue (UAAUAAA) were used to introduce the early stop signal in the exon 3 of *Eprs* gene, resulting in a null allele. For experiments with *Eprs*^+/–^ mice, control mice of the same age and gender from littermates or sibling mating were used. All animal procedures were performed following the National Institutes of Health (NIH) and the University of Rochester Institutional guidelines. Two different models of HF were used: (1) ISO infusion model (20 mg/Kg/day; neural hormonal model). (2) Transverse aortic constriction (TAC; surgical model). *Eprs* conditional knockout (cKO) mouse line *Eprs*_tm1c_B03 was purchased from The Center for Phenogenomics (TCP, Toronto, Canada) in the form of frozen sperms^1^. The *Eprs*^flox/+^ tm1c cKO mouse line was rederived using In Vitro Fertilization (IVF) was performed by the Mouse Genome Editing Resource at URMC. The *Eprs* cKO tm1c mouse line was bred with *Postn*^MCM/+^ mice^2^ to obtain tamoxifen-inducible myofibroblast-specific *Postn*-Cre-driven *Eprs*^flox/+^ tm1d cKO mouse line. We used one single initial dose of 30 µg/g of mouse body weight for tamoxifen (TMX) intraperitoneal injection followed by TMX food for the entire pathological stimulus of TAC surgery. For group size justification, we have performed a power analysis for One-way ANOVA. The mice are randomized for experiments. Animal operations, including ISO infusion, TAC surgery, and echocardiography measurement, were performed blindly by the Microsurgical Core surgeons. Sections and histology analysis were done by the Histology Core.

### Analysis of Translational Activity by Polysome Profiling

Polysome profiling for cytosolic translation was performed as previously described.^20^ Cycloheximide (CHX, 100 µg/ml) was added to the cells for 15 mins before lysis to freeze ribosomes on mRNAs in the elongation phase. Around 10^7^ cells were lysed in TMK lysis buffer (10 mM Tris-HCl pH 7.4, 100 mM KCl, 5 mM MgCl_2_, 1% Triton X-100, 0.5% Deoxycholate, 2 mM DTT) containing 100 µg/ml CHX, 4 U/ml RNase inhibitor (NEB) and proteinase inhibitor cocktail (Roche) on ice for 20 mins. Equal amounts of A_260_ absorbance from each sample were loaded onto a 10-50% sucrose gradient solution and centrifuged at 29,000 rpm for 4 hrs; 22 translation fractions were collected from each sample by Density Gradient Fractionation System (BRANDEL). Based on the UV absorbance curve, the 22 fractions were pooled into 7 samples for total RNA extraction and RT-qPCR analysis, including free mRNP, 40S small ribosome subunit, 60S large ribosome subunit, 80S monosome, light polysomes (di-ribosome, tri-ribosome, etc.), and heavy polysomes (>5 ribosomes). For polysome-Seq, all the fractions were pooled into 3 samples, non-polysome (free mRNP, 40S, 60S subunit), light polysome (monosome, disome, trisome, tetrasome), and heavy polysome (>5 ribosomes). Total RNA was extracted from the same volume of each pooled fraction with Trizol LS (ThermoFisher Scientific), and Renilla luciferase mRNA from *in vitro* transcription was used as RNA spike-in and a loading control for RT-qPCR. The RNA-Seq and polysome-Seq data were uploaded to NCBI GEO database with ID of GSE136838.

### Statistical Analysis

All quantitative data were presented as mean ± SEM and analyzed using Prism 7 software (GraphPad). For a comparison between 2 groups, a Student *t* test was performed. For multiple comparisons among ≥3 groups, 1-way ANOVA was performed. For the counts of proline-rich genes, χ^2^ test was performed. Statistical significance was assumed at a value of *P*≤0.05. Further details and a complete collection of methods are available in the Online Data Supplement.

## Results

### EPRS is upregulated in human and mouse heart failure

To screen for candidate translation factors involved in cardiac fibrosis, we re-analyzed RNA-Seq data of TGF-β (a typical pro-fibrotic cytokine^21^) treated human CFs from CVD patients^4^. We have determined cytoplasmic ARSs as the only significantly induced translation machinery components (Figure S1A) among ribosome proteins and all the translation factors (Figure S1B, Table S1)^4^. Among 20 ARSs, we have identified EPRS as the key ARS involved in cardiac pathogenesis by screening of ARSs induced in TGF-β-activated human CFs^4^, human ARSs with genetic mutations in congenital heart disease^22^, mouse ARSs associated with isoproterenol (ISO)-induced cardiomyopathy by genome-wide association studies (GWAS)^23^, and ISO-induced ARSs in mouse failing hearts^24^ (Figure 1A). Multiple ARS proteins are induced in skeletal muscle of humans during exercise training, which is considered to be a mechanism to support protein synthesis and muscle growth^25^. EPRS is not among this group of ARSs required for physiological muscle hypertrophy (Figure 1A, dotted circle). To validate this observation, we measured EPRS expression in our own cohort of human and mouse heart samples. Our results showed that EPRS was induced at both mRNA and protein levels in failing hearts from heart failure patients, compared to non-failure donor hearts (Figure 1B-D). Also, EPRS was induced at both mRNA and protein levels in the hearts from mice that underwent ISO infusion compared to those with vehicle treatment (Figure 1E and 1F). Consistently, EPRS was also induced at both mRNA and protein levels in the hearts from mice under TAC surgery compared to those with Sham operations (Figure 1G and 1H). All these data indicate that EPRS may play a critical role in cardiac pathogenesis.

**Figure 1.**
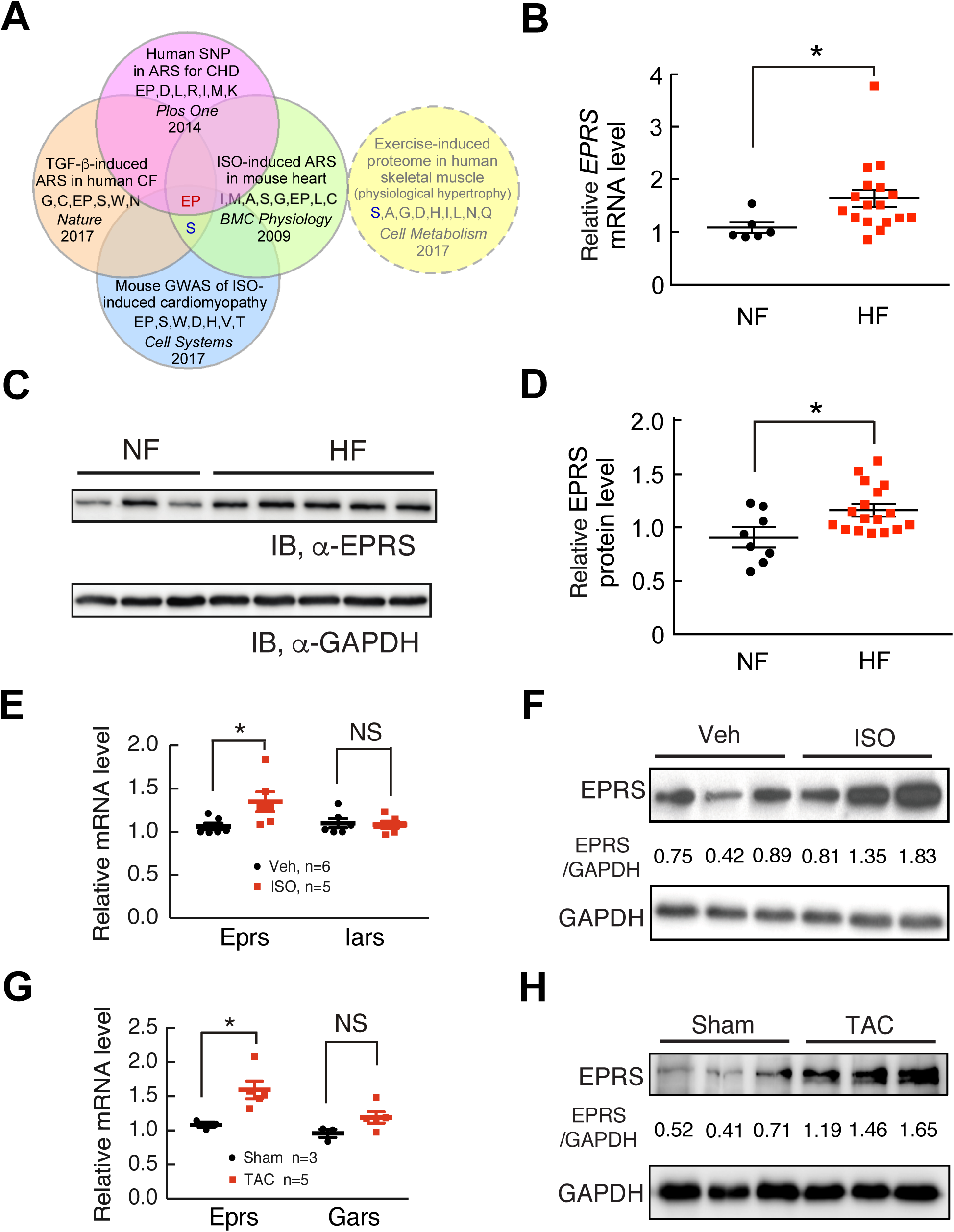
EPRS is upregulated in human and mouse heart failure. **(A)** EPRS is a common hit of upregulated ARSs and ARSs with a genetic association in human and mouse heart disease, but not induced in human physiological muscle hypertrophy. **(B)** *EPRS* mRNA is increased in failing human hearts (n=17) compared to non-failing donor hearts (n=6). 18S rRNA is used as a a loading control. **(C)** EPRS protein is increased in failing human hearts compared to non-failing donor hearts. Representative Western blot results are shown. **(D)** Quantification of Western blot results from all human heart samples indicate elevated expression of EPRS in failing human hearts (n=17) compared to non-failing donor hearts (n=8). GAPDH is used as a loading control for quantification. **(E-F)** EPRS mRNA and protein expression are increased in the hearts from a 4-week ISO infusion induced mouse HF model compared to vehicle infused hearts while *Iars* (Isoleucyl-tRNA synthetase) mRNA stays unchanged. 18S rRNA and GAPDH are used as loading controls for mRNA and protein quantification, respectively. **(G-H)** EPRS mRNA and protein expression are increased in the hearts from 8-week TAC surgery-induced mouse HF model. *Gars* (glycyl-tRNA synthetase) is a negative control ARS in the TAC model. 18S rRNA and GAPDH are used as loading controls for mRNA and protein quantification, respectively. *: p≤0.05, NS: not significant by unpaired two-tailed student t-test.

### Single allele knockout of *Eprs* attenuates cardiac fibrosis under pathogenic stresses

To determine the role of EPRS during cardiac pathogenesis, we first generated *Eprs* global KO mouse using CRISPR technology. The global *Eprs* KO mice were created by introducing two tandem premature stop codons plus frame-shifting adenosine nucleotide in the 3^rd^ exon of *Eprs* gene with protospacer adjacent motif (PAM) mutations (Figure S2A). TA-clone and DNA sequencing confirmed the successful insertion of premature stop codons into the *Eprs* genomic sequence (Figure S2B and S2C). Homozygous global *Eprs* KO mice are embryonic lethal. The heterozygous (het) appeared normal in weight (WT: 26.58±0.44g, n=8; *Eprs*^+/–^: 27.27±0.45g, n=6, male mice at the age of 3-4 months) and fertility (not shown). EPRS expression was reduced by almost half in the heart of *Eprs*^+/–^ mice (Figure S2D). These indicate that *Eprs* is an essential gene, and loss of one allele does not cause developmental defects in mice at baseline.

Given the induction of EPRS expression in human and mouse failing hearts, we tempted to determine the effects of reduced EPRS on the heart during cardiac remodeling *in vivo*. We first used a β-adrenergic receptor agonist ISO to treat mice via osmotic minipump implantation (20 mg/Kg/day) for 4 weeks. The hearts of WT and *Eprs*^+/–^ mice were comparable in size at baseline, as indicated by heart weight/tibia length (HW/TL) ratio (Figure 2A) and wheat germ agglutinin (WGA) staining (Figure 2B). However, *Eprs*^+/–^ mice exhibited a reduced cardiac hypertrophy phenotype compared to WT mice after ISO treatment (7.27±0.31 (Mean±SEM) in *Eprs*^+/+^ to 6.17±0.18 in *Eprs*^+/–^ of HW/TL ratio and 474.8±7.59 μm^2^ in *Eprs*^+/+^ to 348.4±3.96 μm^2^ in *Eprs*^+/–^ of left ventricle (LV) myocyte area) (Figure 2A and 2B). Picrosirius red staining showed that *Eprs*^+/–^ mice had much less cardiac fibrosis (3.17±0.26% to 1.90±0.22%) after ISO treatment than WT mice (Figure 2C).

**Figure 2.**
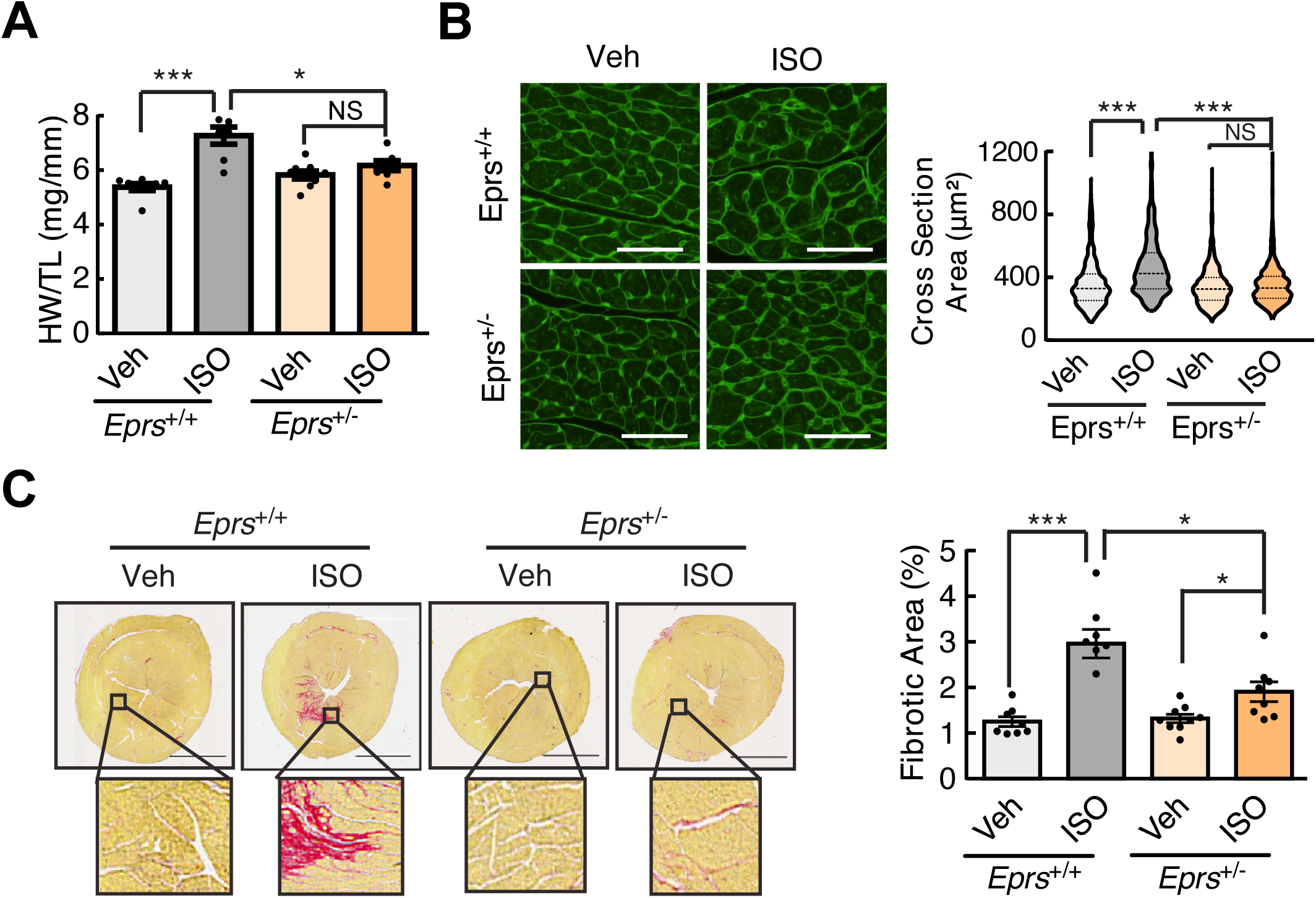
*Eprs* global knockout attenuates cardiac hypertrophy and fibrosis in the ISO infusion model. **(A)** *Eprs*^+/-^ mice show a reduced HW/TL ratio in 4-week ISO infusion mouse HF model compared to WT (*Eprs*^+/+^) mice. n=7-9 for all the groups. **(B)** WGA staining of *Eprs*^+/+^ and *Eprs*^+/-^ mice with ISO treatment. Cross-sectional area (CSA) of CMs was measured and quantified. n = 4-5 hearts per group with 200-300 CMs measured per heart. Scale bar: 50 µM. **(C)** Picrosirius red staining indicates a decreased fibrotic area in the hearts from *Eprs*^+/-^ mice after ISO infusion. Scale bar: 2 mm. n=7-9 for all the groups. *: p≤0.05, **: p≤0.01, ***: p≤0.001 by one-way ANOVA for more than two groups.

We then used a transverse aortic constriction (TAC) surgery, a pressure overload-induced cardiac hypertrophy model, to confirm the role of EPRS in HF. We found that *Eprs*^+/–^ mice exhibited reduced cardiac hypertrophy (9.61±0.57 in *Eprs*^+/+^ to 7.31±0.30 in *Eprs*^+/–^ of HW/TL ratio and 809.1±12.44 µm^2^ in *Eprs*^+/+^ to 355.2±9.12 µm^2^ in *Eprs*^+/–^ of LV myocyte area) compared to WT mice after TAC surgery (Figure 3A and 3B). Cardiac fibrosis was reduced by ∼40% (3.58±0.44% to 2.16±0.28%) in *Eprs*^+/–^ mice compared to WT mice after TAC (Figure 3C). Moreover, the echocardiography showed partially restored cardiac function in *Eprs*^+/–^ mice compared to WT mice after TAC surgery (Figure 3D and 3E, Table S2). In summary, *Eprs*^+/–^ mice exhibited reduced cardiac hypertrophy, fibrosis, and improved cardiac function in HF models under stress conditions.

**Figure 3.**
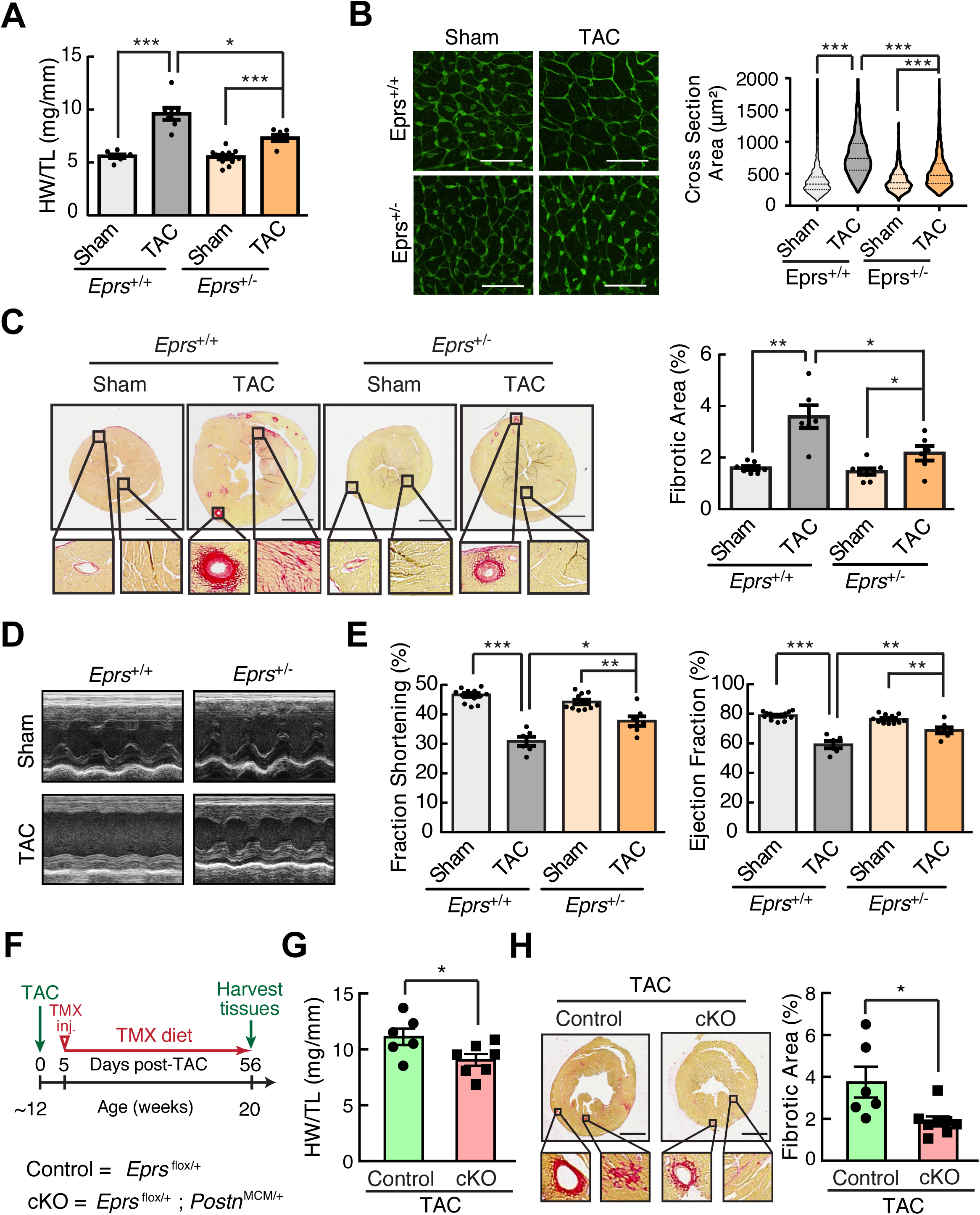
*Eprs* global and myofibroblast-specific conditional knockout reduces cardiac hypertrophy and fibrosis in the TAC surgery model. **(A)** Single allele knockout of *Eprs* reduces HW/TL ratios in TAC surgery-induced mouse HF model. n=7-11 for HW/TL quantification in all the groups. **(B)** WGA staining in *Eprs*^+/+^ and *Eprs*^+/-^ mice with TAC surgery. n = 4-5 hearts per group with 200-300 CMs measured per heart. Scale bar: 50 µM. **(C)** Single allele knockout of *Eprs* reduces cardiac fibrosis in TAC surgery model. n=6-8 for quantification of picrosirius red staining in all the groups. Scale bar: 2 mm. **(D)** Representative echocardiographic images suggest improved cardiac function in *Eprs*^+*/-*^ mice compared to WT mice after TAC surgery. **(E)** Fraction shortening (FS) and ejection fraction (EF) are partially recovered in *Eprs*^+/-^ mice in TAC surgery-induced mouse HF model. n=6-14 for all the groups. **(F)** Schematic of the experimental procedure of TAC surgery in tamoxifen (TMX)-induced myofibroblast-specific *Eprs* conditional knockout (cKO, tm1d) and *Eprs*^flox/+^ mice (control, tm1c). **(G)** HW/TL ratios suggest attenuated hypertrophy in *Eprs* cKO mice (n=7) after TAC surgery compared to control mice (n=6). **(H)** Picrosirius red staining indicates reduced cardiac fibrosis in *Eprs* cKO mice (n=7) compared to control mice (n=6) after TAC surgery. Scale bar: 2 mm. *: p≤0.05, **: p≤0.01, ***: p≤0.001, NS: not significant by one-way ANOVA test.

To test whether haploinsufficiency of EPRS in cardiac myofibroblast reduces cardiac fibrosis, we obtained *Eprs* floxed mice from the International Mouse Phenotyping Consortium and breed with tamoxifen (TMX) inducible Postn-Cre mice (Postn^MCM/+^) to generate myofibroblast-specific TMX-inducible Postn-Cre-driven single allele *Eprs* conditional knockout (cKO) mice (Figure S2E). The *Eprs* WT control (*Eprs*^flox/+^) and *Eprs* cKO (*Eprs*^flox/+^; *Postn*^MCM/+^) mice were subject to TAC surgery and fed with TMX food for 2 months (Figure 3F). We observed significantly reduced HW/TL ratio in *Eprs* cKO mice compared to WT control mice after TAC surgery (Figure 3G). More importantly, fibrotic area was significantly reduced by ∼49% in *Eprs* cKO mice (3.79±0.73% in control to 1.93±0.24% in cKO) compared to control mice indicated by the picrosirius red staining (Figure 3H).

Taken together, all these data suggest that global or myofibroblast-specific single allele KO of *Eprs* attenuates cardiac remodeling under pathogenic stresses and sufficient or elevated expression of EPRS is essential for pathological cardiac remodeling.

### Sufficient EPRS is required for efficient translation of Pro-rich collagen proteins

Pearson correlation analyses in human HF samples indicate that EPRS expression is correlated with collagens, but not ANF and BNP expression (Figure 4A and S3A). This observation suggests that EPRS may directly regulate the expression of collagens in CFs. In primary cultured adult mouse CFs, we found that pro-hypertrophic (ISO and Ang II), as well as pro-fibrotic agonists (TGF-β and IL-11) significantly increased *Eprs* expression by 1.3-2.7 folds (Figure S3B). We confirmed that EPRS expression was increased by ISO at the mRNA and protein levels in mouse CFs due to increased transcription (Figure 4B and 4C). mRNA of Pro-rich (PRR) collagen genes, including COL1A1 and COL3A1, was significantly induced by ISO at the protein level (3-5 folds) but not markedly at the mRNA level (∼1.2-1.5 folds), suggesting a preferential translational regulation of collagen genes by ISO (Figure 4D-F). In addition, ISO stimulus did not significantly alter the mRNA expression of *Acta2* and *Postn*, while TGF-β remarkably increased their expression (Figure S3C). Therefore, we chose ISO treatment for examining the translational regulatory effects of EPRS in CFs. Polysome profiling-RT-qPCR assay showed a significantly increased association of *Col1a1* and *Col3a1* mRNAs with heavy polysomes after ISO treatment and reduced polysome association by Halo exposure, suggesting that ISO-enhanced translation efficiency of collagen transcripts is reversed by Halo (Figure 4G). Furthermore, haploinsufficiency of EPRS in isolated mouse primary CFs resulted in a similar translational reduction of collagen mRNAs (Figure 4H). All these results indicate that a sufficient amount of EPRS is critical to maintaining the efficient translation of proline-rich collagens upon ISO stimulation in cardiac fibroblasts.

**Figure 4.**
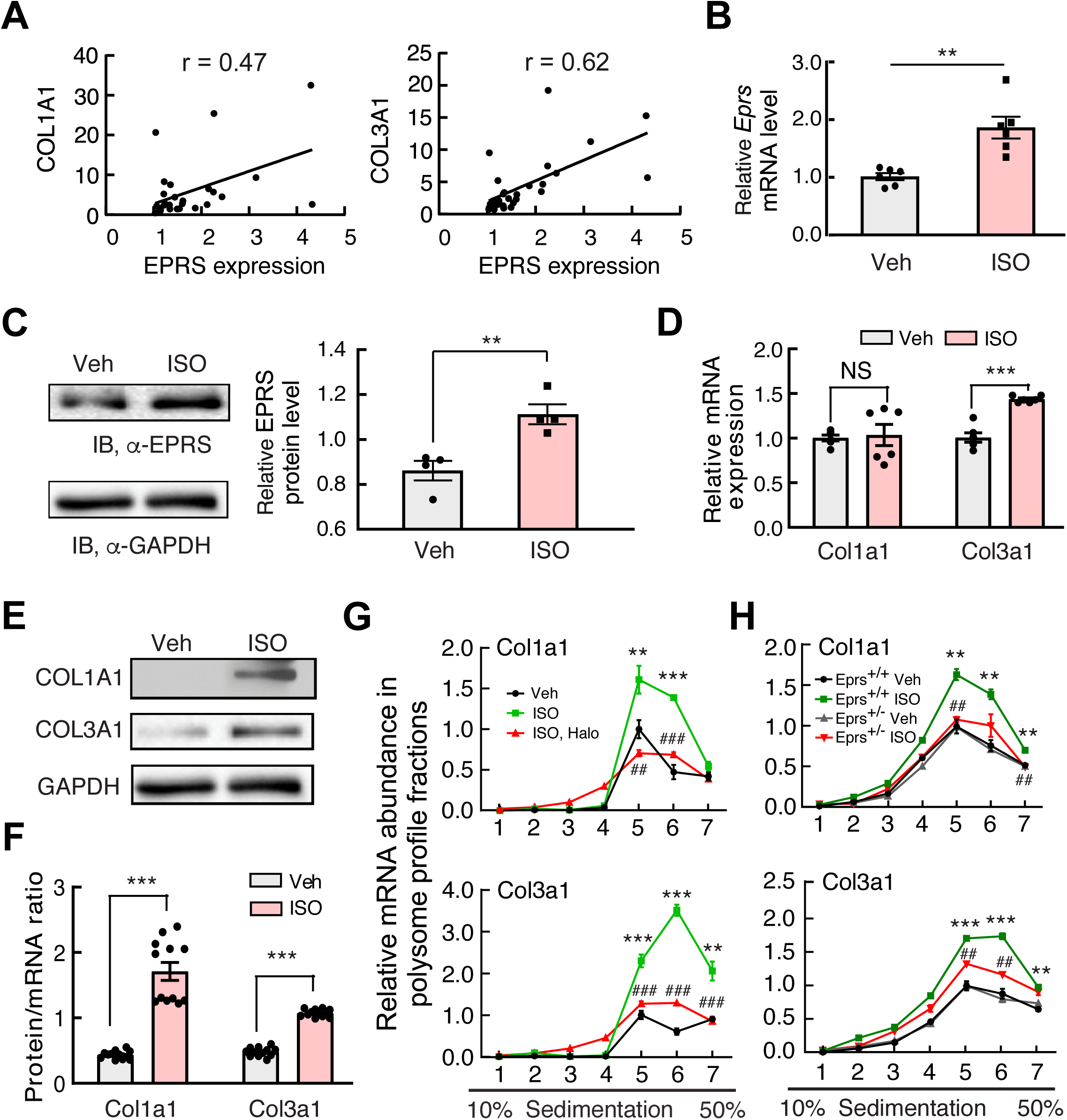
EPRS is required for Pro-rich collagen translation in cardiac fibroblasts. **(A)** EPRS expression is correlated with the expression of collagens in human heart samples (n=23). **(B-C)** ISO induced the expression of EPRS mRNA and protein in primary mouse cardiac fibroblasts (CFs). 18S rRNA and GAPDH are used for loading controls for mRNA (n=6) and protein (n=4) quantification, respectively. **(D)** The expression of collagen mRNAs is slightly increased in ISO-treated primary mouse CFs (n=6). 18S rRNA is used as a loading control. **(E)** Representative Western blot images from two biological replicates indicate that collagen proteins are strongly induced by ISO in mouse primary CFs. **(F)** Protein/mRNA ratio of collagen genes are strongly increased in ISO-treated primary mouse CFs. The protein/mRNA ratio was calculated by pair-wise comparison of quantification of two biological replicates of Western blot in **(E)** with six biological replicates of mRNA expression in **(D)**. **(G)** Polysome-qPCR assay indicates that EPRS inhibition by Halo reverses ISO induced polysome association with collagen mRNAs in primary mouse CFs. Twenty-two fractions were pooled into 7 samples for total RNA extraction and RT-qPCR analysis, including free mRNP (fraction 1), 40S small ribosome subunit (fraction 2), 60S large ribosome subunit (fraction 3), 80S monosome (fraction 4), light polysomes (di-ribosome, tri-ribosome; fraction 5 and 6), and heavy polysomes (>5 ribosomes; fraction 7). *In vitro* transcribed Renilla luciferase mRNA was used as a spike-in control for normalization. *: Veh vs. ISO; #: ISO vs. ISO, Halo. **, ##: ≤0.01; ***, ###: ≤0.001. **(H)** Polysome associated collagen mRNAs are reduced in *Eprs*^+/-^ CFs compared to WT (*Eprs*^+/+^) CFs after ISO treatment. *: *Eprs*^+/+^, ISO vs. Veh; #: *Eprs*^+/-^, ISO vs. *Eprs*^+/+^, ISO. **, ##: ≤0.01; ***, ###: ≤0.001. *: p≤0.05, **, ##: p≤0.01, ***, ###: p≤0.001, NS: not significant by student t-test for two groups and one-way ANOVA for more than two groups.

### Global identification of novel Pro-rich proteins as preferential EPRS targets

As we have shown that collagens, as exemplary PRR genes, are translationally regulated by EPRS, we ought to determine whether EPRS regulates additional preferential mRNA targets rather than all the transcripts in fibroblasts. To address this question, we choose NIH/3T3 mouse fibroblasts as a cell model system to perform the high-throughput screening based on high reproducibility and its ability to recapitulate gene regulatory events from primary CFs. We treated fibroblasts with vehicle or low-dose Halo (100 nM) and performed RNA-Seq (transcriptome profiling) and polysome profiling-Seq (translatome profiling). We identified novel PRR gene pathways that are preferentially regulated by EPRS via enhanced translation elongation at Pro-rich codons and antagonized by Halo (Figure 5A). We categorized differentially regulated genes into 4 groups (Area 1-4) based on their change at the steady-state RNA level and the ratio between a heavy (or light) polysome fraction and a non-polysome fraction (translation efficiency, TE) (Figure 5B and S4A, Table S3 and S4).

**Figure 5.**
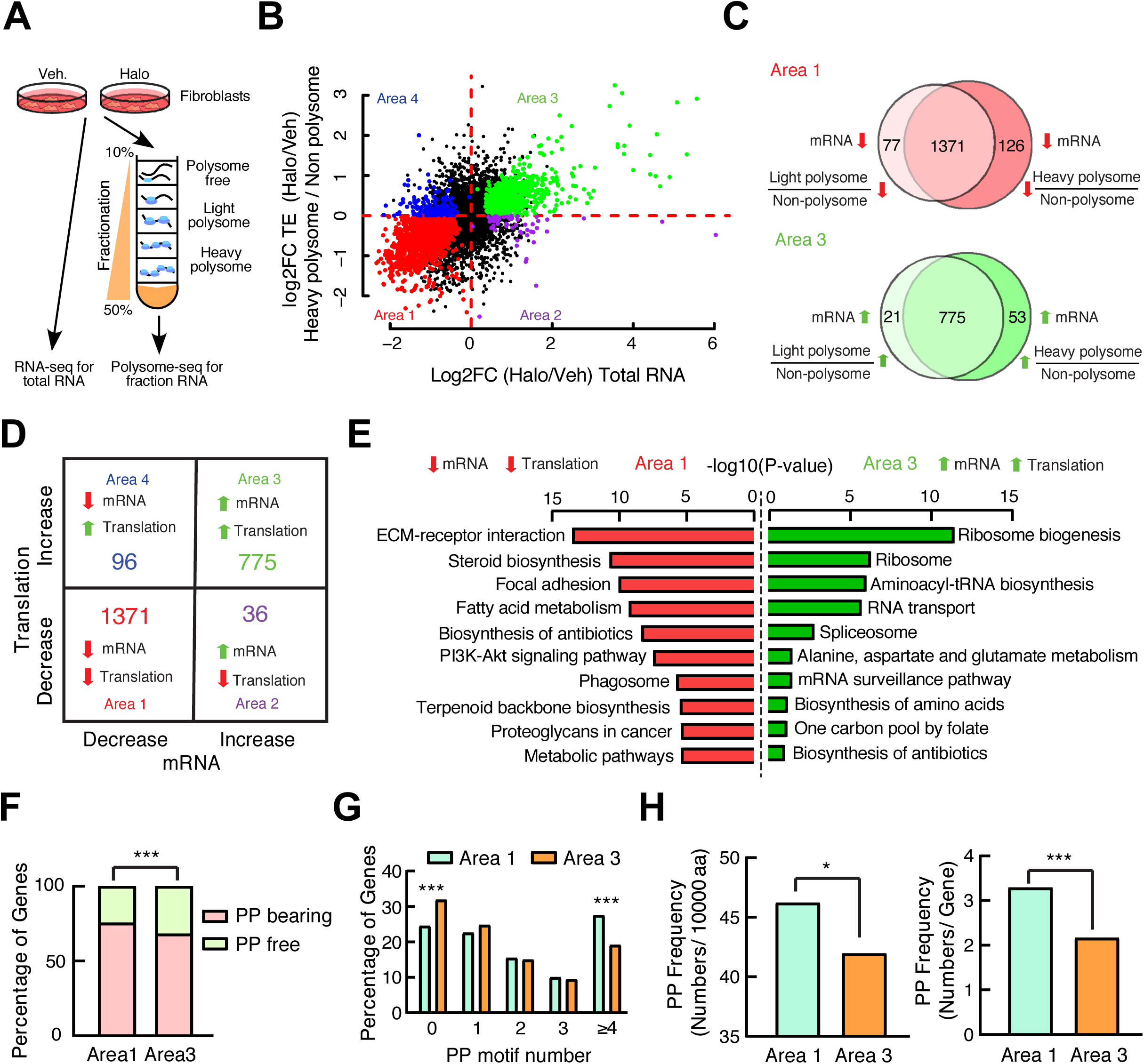
Pro-rich genes are preferential translational targets of EPRS. **(A)** Schematic procedure of RNA-Seq and polysome profiling-Seq (polysome-Seq) in Halo treated fibroblasts. Polysome-free fractions include free mRNP, 40S ribosome subunit, 60S ribosome subunit. Light polysome fractions include 80S monosome, disome, trisome, and 4 ribosomes. Heavy polysome fractions include polysome fractions with ≥5 ribosomes. **(B)** Differentially expressed genes identified by RNA-Seq and polysome-Seq are indicated by dot plot. Translation efficiency (TE) is indicated by the ratio of heavy polysome and polysome free fraction. The colored dots indicate statistically significantly changed genes in either RNA-Seq or polysome-Seq. The genes with p≤0.05 in either of three different groups (ratio of Halo vs. Veh. treated samples for total RNA, non polysome, and heavy polysome) were considered as significantly changed genes. All significantly changed genes were divided into four areas based on log2FC of total mRNAs and heavy polysome mRNAs. Data were submitted to GEO database (GSE136838). **(C)** Translational dysregulated genes are indicated by overlapping changed genes in the same area of heavy polysome **(B)** and light polysome (Figure S4A). Genes decreased (Area 1) or increased (Area 3) at translation, and steady-state mRNA levels are shown. **(D)** Majority of genes show a synergistic change at mRNA and translational levels after EPRS inhibition by Halo. The number of genes is shown with changes at both translation efficiency (the ratio of heavy/light polysome to polysome free fraction) and steady-state mRNA levels in all 4 areas. **(E)** KEGG signaling pathway analyses indicate that ECM receptor interaction and ribosome biogenesis are top enriched pathways in Area 1 and Area 3, respectively. **(F)** PP motif analyses suggest that there are more PP motif-bearing genes in the translationally decreased gene cluster (Area 1) compared to the translationally increased gene cluster (Area 3). **(G)** Distribution of genes according to the number of PP motifs (0, 1, 2, 3, ≥4) indicate that the genes in Area 1 contain more PP motifs compared to genes in Area 3. **(H)** Frequency of PP motif normalized by protein length or gene number is significantly higher in Area 1 compared to Area 3. *: p≤0.05, ***: p≤0.001 by χ^2^ test for gene number counts and student t-test for quantification of PP motif frequency.

Considering the translation active state in both light and heavy polysome fractions, we redefined the 4 groups by overlapping genes from heavy and light polysome associated transcripts (Figure 5C). In the overlapped region of the Venn diagram, 1371 genes were downregulated at translation and steady-state mRNA levels (Area 1), and 775 genes were upregulated at both levels (Area 3). This finding supports the hypothesis that only a small cluster of genes was preferential translational targets of EPRS. Furthermore, most of the significantly changed genes were located in Area 1 and 3 (Figure 5D, and S4B). This observation indicates a coordinated regulatory effect between steady-state mRNA level and translation efficiency under the EPRS inhibitory condition. Kyoto Encyclopedia of Genes and Genomes (KEGG) analysis of the genes in Area 1 (reduced at RNA and TE levels) using DAVID software revealed multiple pro-fibrotic pathways, including ECM-receptor interaction (11 collagen genes such as Col1a1/1a2/3a1/4a6/5a1/6a1, 4 integrin genes Itga7/a8/b3/b4, etc.) (Figure S4C) and proteoglycans (Fgfr1, Pdgfra/rb, Tgfb2/b3, Tgfbr1/r2, Thbs1/3, etc.) (Figure 5E and Table S5). In contrast, the genes in Area 3 (induced at RNA and TE levels) are enriched in general translation factors, including ribosome biogenesis, ribosome (17 cytosolic and 6 mitochondrial ribosome proteins), aminoacyl-tRNA biosynthesis (16 ARSs including *Eprs*) (Figure S4D), and amino acid biosynthesis (Figure 5E), which suggests an activation of a compensatory amino acid starvation response upon EPRS inhibition^12, 15^.

Previous studies demonstrate that the formation of peptide bond inside iterated proline or proline with other amino acids is much slower than that of the non-proline-bearing peptide bond, and thus Pro-Pro (PP) motifs function as ribosome pausing sites during translation elongation^26, 27^. This is an evolutionarily conserved mechanism from bacteria to human and is also unique to Pro codon among all genetic codons ^28, 29^. Therefore, we hypothesize that the translation of genes bearing more PP motifs is more sensitive to mild EPRS inhibition than that of non-PP-bearing genes. Consistent with this hypothesis, the Area 1 gene cluster contains a larger number of PP motif-bearing genes compared to Area 3 (Figure 5F, Table S6). The percentile of genes bearing more than 4 PP motifs in Area 1 is much higher than that in Area 3 (Figure 5G, Table S6). The overall frequency of PP motifs in Area 1 is also significantly higher than that in Area 3 as indicated by the number of PP motifs per 10,000 amino acids or per transcript (Figure 5H, Table S6). Major cardiac fibrosis marker genes in Area 1 gene cluster, such as *Col3a1* and *Col1a1* (and other collagen genes), contain highly enriched Pro and Gly genetic codons due to abundant Pro-Pro-Gly (PPG) motifs in comparison to overall codon composition in the mouse genome (Figure S5A and S5B). Polysome profiling followed by RT-qPCR confirmed that collagen transcripts were reduced in light polysome (fraction 5) and heavy polysome fractions (fraction 6 and 7), suggesting inhibition of EPRS by Halo causes ribosome stalling at PRR codons and decreased translation elongation of collagen mRNAs (Figure S5C and S5D). In addition, gene regulation changed in the opposite direction at the mRNA and TE levels in Area 2 and 4 in which a small number of genes are located (Figure S5E and S5F, Table S4). Taken together, all these results suggest that PP motif bearing genes are preferential targets of EPRS, and mild inhibition of EPRS by Halo leads to inefficient translation of these Pro codon rich genes.

### SULF1 and LTBP2 are novel EPRS downstream effectors

To screen for specific novel downstream effectors of EPRS as potential drug target genes for anti-fibrotic treatment, we performed a high-throughput bioinformatic screening by integrating genomic, transcriptomic, translatomic, and proteomic analyses (Figure 6A). We first performed in silico bioinformatic analyses to uncover conserved PRR genes that bear PP genetic codon motifs across the reviewed human and mouse proteins from Uniprot proteomic sequences^30^ at the genome-wide scale (Table S7), including PPG-rich ECM genes such as collagens among many Pro-rich proteins. Next, we overlapped PP motif bearing genes with genes that were downregulated in our transcriptomic and translatomic analyses (Figure 6B) and proteins that were reduced at the steady-state level in human CFs upon low-dose Halo treatment in the published quantitative proteomic data^15^ (Figure S6A, Table S8). As we expected, 8 collagen genes (i.e., COL1A1/1A2/3A1/5A1/ 6A1/6A2/6A3/12A1) were translationally reduced in mouse fibroblasts, and their steady-state protein level was also decreased in human CFs after Halo treatment. Another 6 collagen genes (COL4A6/5A3/11A1/16A1/27A1/28A1) were reduced at the translation efficiency level in mouse fibroblasts, and 5 collagen genes (COL4A1/5A2/7A1/14A1/18A1) were reduced at the protein level in human CFs. Intriguingly, a recent new cardiac fibrosis marker gene cytoskeleton associated protein 4 (CKAP4) was found as a PRR gene downregulated in our polysome-Seq analysis (Table S4 and S8). We also observed that GTPBP2 (a ribosome rescue factor for dissembling stalled ribosome)^31^ (Figure S6A) and EIF5A (an elongation factor)^26, 27^ are both induced by Halo treatment (Table S4). The induction of GTPBP2 indicates enhanced recycling of stalled ribosomes during Halo exposure. EIF5A is required for ribosome readthrough of consecutive Pro codons in PRR genes, and thus its increase serves as another compensatory response upon Halo-driven inhibition of Pro codon decoding^26, 27^. These results suggest a previously unappreciated cellular adaptive response to promote the recycling of stalled ribosomes caused by EPRS inhibition and facilitate the readthrough of PP motifs. By contrast, most of the house-keeping genes are not decreased at translational efficiency or steady-state protein level, includes all histone genes, majority of DNA polymerase complex genes, RNA polymerase complex genes, 38 ARSs, all heterogeneous nuclear ribonucleoproteins, GAPDH, vimentin, β-actin among many others. These observations further support the idea of EPRS-mediated selective regulation of PRR mRNA translation.

**Figure 6.**
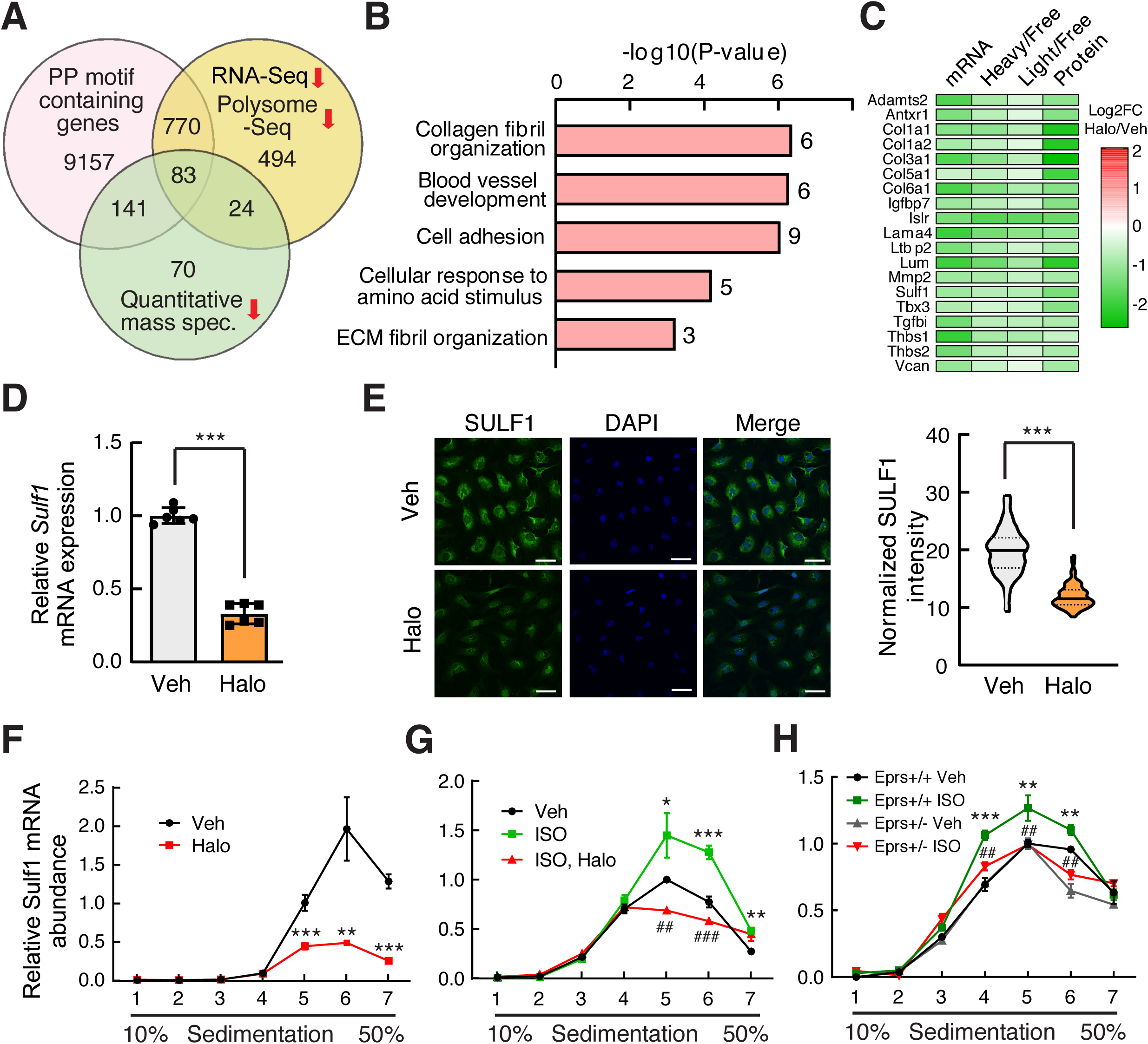
SULF1 is a novel EPRS downstream target. **(A)** Integrated genomic, transcriptomic, translatomic, and proteomic^15^ analyses identify 83 PP motif-containing genes as downstream targets of EPRS. **(B)** Gene Ontology analyses are performed for 83 Halo-downregulated PRR genes from **(A)**. Top 5 enriched pathways from the GO analysis are shown. **(C)** The log2FC change of mRNA, translation, and steady-state protein levels of all the genes from **(B)** are shown by heatmap. Steady-state protein level changes were obtained from a published database^15^. **(D-E)** *Sulf1* mRNA **(D)** and protein **(E)** expression are reduced by in EPRS inhibition in mouse primary CFs. 18S rRNA is used as a loading control for mRNA level. Scale bar: 50 µM. **(F)** Polysome associated *Sulf1* mRNA is reduced in Halo-treated primary CFs. **(G)** EPRS inhibition reverses ISO-induced polysome association of *Sulf1* mRNA in CFs. *: Veh vs. ISO; #: ISO vs. ISO, Halo. **(H)** Polysome associated *Sulf1* mRNA is reduced in *Eprs*^+/-^ CFs compared to WT (*Eprs*^+/+^) CFs after ISO treatment. *: *Eprs*^+/+^, ISO vs. Veh; #: *Eprs*^+/-^, ISO vs. *Eprs*^+/+^, ISO. *: p≤0.05, **, ##: p≤0.01, ***, ###: p≤0.001, NS: not significant by student t-test for two groups and one-way ANOVA for more than two groups.

By overlapping all three datasets, a group of 83 PP motif-containing genes was shown to be downregulated at translational and steady-state protein levels (Figure 6A, Table S8). Gene Ontology (GO) analyses revealed top 5 pathways, including collagen fibril organization (COL1A1, COL1A2, COL3A1, COL5A1, ADAMTS2, LUM), cell adhesion (SULF1, TGFBI, VCAN, THBS1, THBS2, LAMA4, IGFBP7, LSLR), and extracellular fibril organization (COL3A1, COL5A1, LTBP2) among others, many of which are involved in cardiac pathological remodeling (Figure 6B and Table S9). The steady-state mRNA expression, translation efficiency, and steady-state protein expression are reduced for the major EPRS preferential target genes from the top 5 GO pathways (Figure 6C). Among these EPRS targets, LTBP2 (latent TGF-β-binding protein 2) has been recently discovered as an ISO-induced fibrosis marker protein^32^ and shown in our screen as downregulated at steady-state mRNA and polysome associated mRNA levels in Halo-treated mouse fibroblasts (Figure S6B). Mouse LTBP2 protein bears 17 PRR motifs (human LTBP2 has 20 PRR motifs) such as PPP and PPG (supplemental sequence information). The polysome associated *Ltbp2* mRNA was significantly reduced by Halo treatment and *Eprs* haploinsufficiency in primary mouse adult CFs (Figure S6C-E).

We have a particular interest in SULF1 which is an endosulfatase that selectively removes 6-O-sulfate from heparin sulfate proteoglycans (HSPGs) and regulates signaling transduction of multiple growth factors^33, 34^, and could be considered as a pharmacological target for anti-fibrosis drug screening based on its enzymatic activity. SULF1 contains PRR codons such as the PPG motif in both human and mouse (plus PPD and PPT motifs in the human gene and PPR motif in the mouse gene) and was downregulated at mRNA, translation, and steady-state protein levels (Figure 6C). We confirmed that SULF1 mRNA and protein expression were downregulated by Halo in mouse CFs (Figure 6D and 6E). Polysome profiling-RT-qPCR was performed to confirm downregulation of SULF1 at the translational level. Halo treatment reduced the polysome association of *Sulf1* mRNA (Figure 6F) and attenuated ISO-induced *Sulf1* mRNA translation (Figure 6G). Moreover, the translation efficiency of *Sulf1* mRNA was dramatically reduced in isolated primary adult CFs from *Eprs*^+/-^ mice compared to WT mice under ISO treatment (Figure 6H). In summary, we validated LTBP2 and SULF1 as novel authentic EPRS-regulated, Halo-responsive Pro codon rich genes.

### SULF1 is a myofibroblast marker and required for cardiac fibroblast activation

As we have shown that SULF1 is a preferential downstream target of EPRS, we ought to examine the expression of SULF1 during cardiac fibrosis and heart failure. By screening a high-throughput microarray-based gene expression database^35^ of human heart failure, we discovered that *SULF1* mRNA expression was significantly induced in dilated cardiomyopathy (DCM) and ischemic heart failure (IHF) patients compared to non-failing donor hearts (Figure S7A). We confirmed this phenomenon using our validation cohort of human HF samples (Figure 7A). In our human heart samples, SULF1 expression is positively correlated (r = 0.58) with the expression of EPRS (Figure S7B), which provides another piece of evidence for SULF1 acting as a downstream effector of EPRS. The positive correlation of SULF1 with COL1A1 (r = 0.60) and COL3A1 (r = 0.67) indicates a potential functional correlation of SULF1 with CF activation (Figure S7C-D). Indeed, SULF1 is highly expressed in α-SMA positive myofibroblasts in failing human hearts (Figure S7E-G) and in the heart of the TAC-induced mouse HF model (Figure S7H and S7I). These data indicate SULF1 as a myofibroblast marker gene during cardiac pathological remodeling and heart failure.

**Figure 7.**
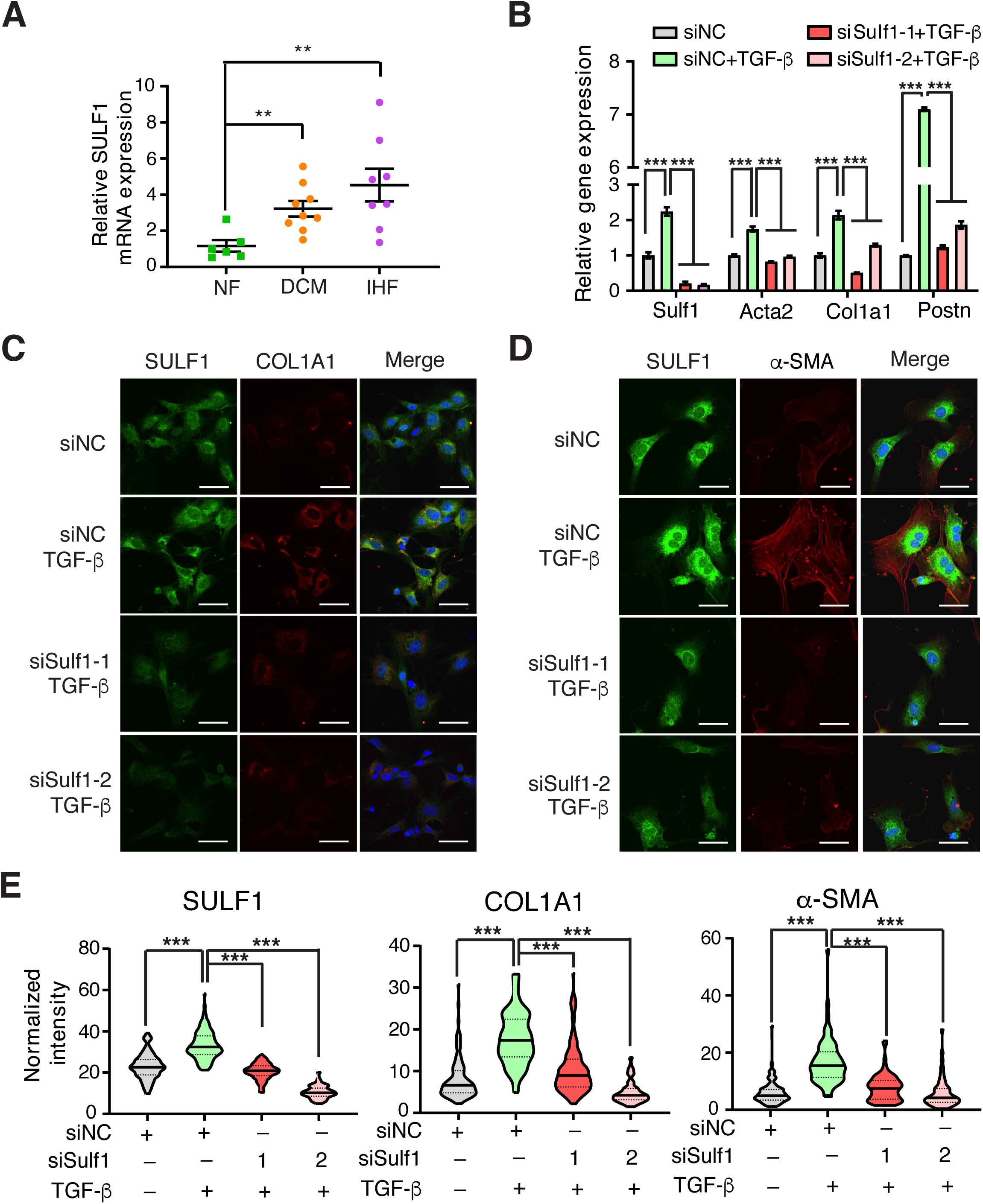
SULF1 is required for cardiac fibroblast activation and collagen deposition. **(A)** RT-qPCR confirms increased SULF1 expression in the hearts of DCM (n=9) and IHF (n=8) patients compared to non-failing donor heart tissues (n=6) in a validation cohort of human samples. 18S rRNA is used as a loading control for mRNA measurement. **(B)** Knockdown of SULF1 attenuates TGF-β-induced cardiac fibroblast activation as indicated by reduced expression of fibroblast activation marker genes. 18S rRNA is used as a loading control. **(C-D)** Immunofluorescence staining shows reduced protein expression of COL1A1 and α-SMA by knockdown of SULF1 in TGF-β-treated primary CFs. Scale bar: 40 µm. **(E)** Quantitation of SULF1, COL1A1, and α-SMA protein expression in immunofluorescence staining in **(C)** and **(D)**. ***: p≤0.001 by student t-test for two groups and one-way ANOVA for more than two groups.

Given the positive correlation between SULF1 and collagens, *Sulf1* mRNA was dominantly expressed in primary mouse CFs rather than CMs (not shown). Next, we examined the effects of knockdown of SULF1 on myofibroblast activation. We transfected mouse CFs with two SULF1-specific siRNAs, and both siRNAs significantly reduced *Sulf1* mRNA expression and TGF-β induced myofibroblast activation marker gene expression (Figure 7B). Meanwhile, immunofluorescence imaging results indicate that reduction of Sulf1 remarkably inhibited TGF-β induced CF activation and collagen deposition (Figure 7C-E). Taken together, our results suggest SULF1 is required for cardiac fibroblast activation and can be used as a myofibroblast marker during cardiac fibrosis and heart failure.

## Discussion

This work is a pioneering investigation of translational control mechanisms of pro-fibrotic gene expression in cardiac fibrosis (Figure 8). In this study, we found that the alteration in EPRS expression in CFs is a conserved aspect of pathological cardiac remodeling in human HF patients as well as in HF mouse models. Using a global and myofibroblast-specific *Eprs* conditional knockout mouse models, we demonstrated that reducing EPRS antagonizes pathological cardiac fibrosis *in vivo*. Mechanistically, we identified a subset of novel preferential translational target genes of EPRS and discovered that EPRS regulates translation of PRR transcripts, which construct most of the ECM and secretory signaling molecules. Among those targets, we identified several novel PRR genes that have not been well studied in the heart (e.g., LTBP2, SULF1, etc.). These genes may serve as novel anti-fibrotic targets for developing therapeutic approaches to treat HF. This study is the first demonstration of the *in vivo* function of EPRS in cardiac remodeling and provides a strong rationale for using translation factors (e.g., EPRS) or their downstream effectors (e.g., SULF1) as novel therapeutic targets against cardiac fibrosis.

**Figure 8.**
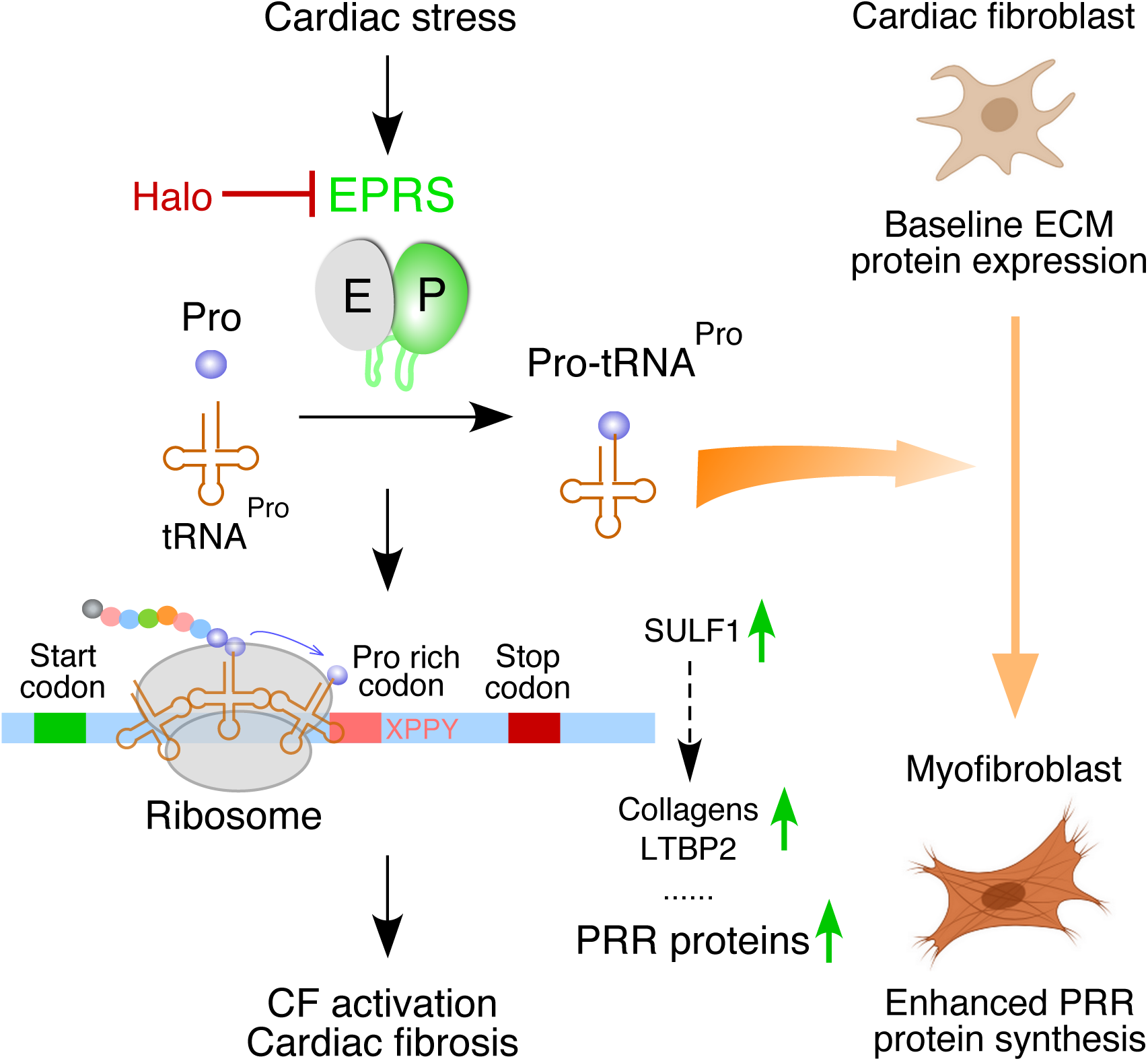
Schematic model of EPRS-mediated translational activation of PRR proteins during cardiac fibrosis.

Two recent reports showed that *EPRS* mRNA is significantly induced as an early response gene at the transcriptional level after 24 hr TGF-β stimulation (Figure S8)^4, 5^. As expected, mRNA expression of well-established fibrosis marker genes is also upregulated, such as ACTA2, POSTN, and collagens. We also observed in their dataset induction in multiple recently reported novel fibrosis marker genes, including LTBP2^32^ and CKAP4^36^. However, translation of Pro-rich pro-fibrotic mRNAs is markedly repressed based on the DeltaTE values (translation efficiency calculated by the ratio of Ribo-Seq vs. RNA-Seq signals) such as COL1A1, COL1A2, COL3A1, LTBP2, and CKAP4 at 24 hr time point (Figure S8 and supplemental sequence information)^5^. In contrast, translation is increased for Postn (contain no PRR motif) and not affected for α-SMA (contain a single PP motif) (Figure S8). These observations indicate that translation is a limiting step for efficient synthesis of Pro-rich (not non-PRR) pro-fibrotic proteins in the acute phase of TGF-β stimulation (24 hr TGF-β exposure) despite a high level of transcription. As a consequence, translation efficiency cannot match with the transcriptional surge. Therefore, TGF-β-induced EPRS may be required for efficient translation of Pro-rich genes. IL-11 has been shown as a most highly induced transcriptional effector downstream of TGF-β signaling to activate translation of pro-fibrotic mRNAs^4^. But the mechanism underlying translational control is still unknown. We discovered that EPRS is an integral node downstream of multiple pro-hypertrophic and pro-fibrotic stimuli in murine hearts. EPRS is induced by ISO infusion and TAC surgery *in vivo* as well as by stimulation of ISO, Ang II, TGF-β or IL-11 *in vitro*. Importantly, one study also showed that *Eprs* mRNA expression is induced by 1.29-fold in C57BL/6J and 1.42-fold in C3H/HeJ mouse strains on day 14 after ISO infusion^32^. Taken together, we believe that EPRS is one of the major regulators involved in TGF-β-, IL-11-, and ISO-driven translational activation of ECM protein synthesis in cardiac fibroblasts.

Based on our reanalysis of an existing database (Figure S1A and Table S1)^4, 37^, we found that gene expression change of a group of cytoplasmic ARSs is involved in cellular identity conversion of CF. Upregulation of nine cytoplasmic ARSs occurs in TGF-β-activated CF-to-myofibroblast differentiation, including EPRS, GARS, CARS, SARS, NARS, TARS, MARS, IARS, and WARS. Intriguingly, GMT (Gata4, Mef2c, Tbx5)-driven CF-to-CM trans-differentiation requires downregulation of almost completely overlapped members of ARSs^37^ as found in CF-to-myofibroblast transition, including the same nine ARSs plus LARS, RARS, and KARS. The GO analysis in this report^37^ has shown that ARS and ECM gene pathways are among the top enriched in a gene cluster that is unaltered in gene expression during CM-to-pre-iCM transition and drops during pre-iCM-to-iCM maturation. Among all altered ARSs, EPRS is dramatically reduced during this process, thereby possibly contributing to the silencing of collagen expression during CF-to-CM conversion. These phenomena suggest that the translation state varies in different cardiac cell types ranging from CM, CF, and myofibroblast. Adult CMs do not normally express ECM proteins, and myofibroblast cells express abundant ECM proteins, while CFs express a moderate amount. Therefore, fine-tuning of expression of ARSs, especially EPRS, may account for proper expression of ECM genes at the translational level across various cardiac cell types.

We demonstrate that genetic knockout of one allele of *Eprs* gene can effectively reduce cardiac remodeling and fibrosis (Figure 2 and 3). Myofibroblast-specific conditional knockout of *Eprs* further confirms that fibroblast-derived EPRS contributes significantly to the cardiac fibrosis under cardiac stress conditions (Figure 2 and 3). These genetic manipulations mimic the mild translational inhibition using the (E)PRS-specific inhibitor Halo in multiple heart failure disease mouse models^15^. This new therapeutic approach targeting an evolutionary conserved, ubiquitous house-keeping translation factor can be used to treat cardiac fibrosis of multiple etiologies and is generalizable to antagonize fibrosis from various organs^38^. Recent studies have shown that EPRS directly forms a complex with TGF-β1 receptor, Janus kinases, and STAT6 and regulate ECM expression via activation of TGF-β1 signaling in lung and liver^18, 19^. This may suggest that EPRS may synergistically regulate the expression of Pro-rich ECM at both transcriptional and translational levels during cardiac fibrosis.

By using the (E)PRS inhibitor Halo, we have discovered that the EPRS-PRR axis acts as a novel translational control pathway to regulate pro-fibrotic Pro-rich mRNA translation and cardiac fibrosis (Figure 8). Pro is unique among all 20 amino acids because it forms a cyclic structure between the a-amine group and the side chain. This specific structure dramatically slows down the ribosome decoding of two consecutive Pro codons. This unique feature of Pro probably explains why Halo specifically inhibits PRR mRNA translation, while preserves global mRNA translation by inducing expression of house-keeping genes such as ARSs and ribosome proteins (Figure 5E). Myc, a positive transcriptional regulator of ribosome biogenesis^39^, is also upregulated by Halo (Figure 5B, Table S4). This may partly explain the increased expression of ribosomal genes. In evolution, Pro-rich protein genes are largely introduced in the genome of multi-cellular organisms in comparison to single cellular species during evolution^29^. EF-P (Elongation factor P in prokaryotes)/eIF5A (Elongation factor 5 in eukaryotes) protein is introduced to facilitate efficient peptide bond formation for Pro dipeptidyl motifs^26, 27, 29^ and its upregulation by Halo treatment represents an adaptive response to facilitate ribosome readthrough of PP motifs. Majority of Pro-rich proteins are transmembrane or secretory proteins since the Pro-rich peptides usually play important roles in penetrating plasma membrane for transmembrane localization or secretion^40, 41^. Many of these proteins are either ECM or ligands/receptors for cellular signaling which promote cardiac fibrosis. In another aspect, Postn (contain no PRR motif) was significantly downregulated at the polysome associated mRNA level (Table S4), suggesting that EPRS-independent translational control mechanisms exist and may be affected by Halo indirectly such as the regulation by other RNA-binding proteins^5^. Also, we cannot exclude the possibility of some Pro-rich proteins acting as cardioprotective factors, and thus overall beneficial effects from EPRS inhibition may be from the combined net outcome. Besides the EPRS canonical aminoacylation function, previous studies have shown that EPRS exerts a noncanonical function in translational silencing of inflammation-related mRNAs such as *VEGFA* in monocyte/Mφ as an innate immune response^8, 42–45^. It remains a question whether EPRS from immune cells plays a role in regulating inflammatory response via either the canonical or noncanonical activities.

During the PRR gene screen, we have identified 83 PRR genes besides collagens (Table S8 and S9), which are downregulated by Halo at the posttranscriptional level and may play critical roles in cardiac fibrosis. Induced EPRS can increase the level of Pro-tRNA^Pro^, a building block for the synthesis of Pro-rich proteins (i.e., collagens, LTBP2, etc.). We also observed that translationally reduced mRNAs by Halo tend to have reduced steady-state mRNA level (Figure 5D and S4B). Previous studies have shown that optimal genetic codon composition and translation capacity of mRNAs in human cells significantly influence baseline mRNA stability^46, 47^. We assume that translational repression of PRR mRNA is coupled with enhanced mRNA decay due to reduced translation elongation rate at PP motifs and diminished ribosome-mediated protection of the transcripts during EPRS inhibition. Based on our data, we concluded that cardiac stress-induced EPRS promotes cardiac fibrosis via increased Pro-tRNA^Pro^ and enhanced stabilization and translation of PRR pro-fibrotic mRNAs in cardiac fibroblasts.

Among EPRS target PRR genes, we discovered that SULF1 could serve as a biomarker for cardiac fibrosis. Genetic knockdown of SULF1 by siRNA inhibits activation of myofibroblasts stimulated by TGF-β. SULF1 and SULF2 are two redundant homolog enzymes of heparan sulfate 6-*O*-endosulfatases^48^. SULF1 catalyzes the removal of 6-*O*-sulfate from heparan sulfate chains and modulates the function of multiple growth factors^34, 49, 50^. Genetic knockout of Sulf1 in mouse genome did not cause any developmental, behavioral, histological, and aging-related phenotypes but knockout of Sulf2 led to reduced body weight^48, 49^. A previous study reported that SULF1 was induced by TGF-β in both cultured human lung fibroblasts *in vitro* and in murine lungs *in vivo*, and siRNA-mediated knockdown of SULF1 promotes α-SMA expression and TGF-β signaling ^51^, which is inconsistent with our results here. This discrepancy may suggest the function of SULF1 in cardiac fibroblasts could be different from that in lung fibroblasts. A recent report showed that SULF1 is predominately expressed in endothelial cells and cardiac fibroblasts in the murine heart and overexpressed SULF1 induces pro-fibrotic gene expression signature in mouse embryonic fibroblasts^52^, which supported our findings to some extent. However, global knockout of SULF1 attenuates angiogenesis and cardiac repair after the ischemic injury without affecting interstitial fibrosis in the non-infarcted regions^52^. Obviously, SULF2, most highly expressed in monocytes and macrophages, could be secreted by these myeloid cells and compensate for the loss of SULF1 function *in vivo*. In our study, we showed the essentiality of fibroblast-derived SULF1 in myofibroblast activation in primary CF culture. Therefore, it is worthwhile to further elucidate the pathophysiological function of endothelial cell and cardiac fibroblast derived SULF1 in cell type specific KO mouse models *in vivo.* These studies will provide insight into whether cell type specific loss of SULF1 is beneficial or detrimental during cardiac remodeling, especially in non-ischemic heart disease, in which less inflammation and immune response are activated and less SULF2 will be expressed. Our studies suggest that a better understanding of the mechanisms of translational control in the cardiac fibrosis has broad implications for developing novel treatment strategies to combat CVD by either targeting a specific translation factor or its downstream effectors.

---

**Figure S1.**
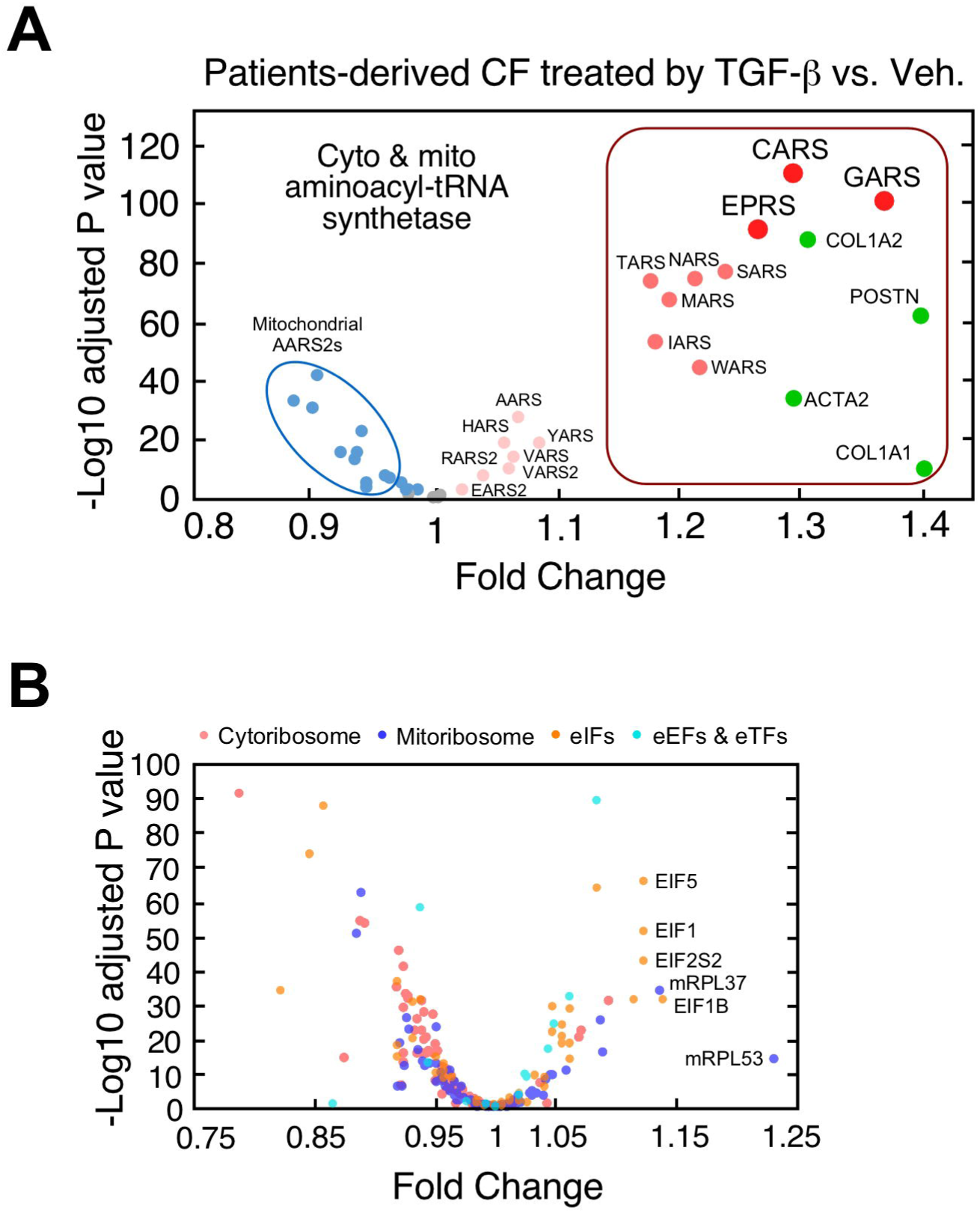
Upregulation of EPRS in TGF-β activated human cardiac fibroblasts. **(A)** Transcriptional activation of cytosolic ARSs, including EPRS in TGF-β stimulated human CFs. **(B)** No remarkable transcriptional activation of other translation factors in TGF-β stimulated human CFs. eIF: eukaryotic initiation factor; eEF: eukaryotic elongation factor; eTF: eukaryotic termination factor.

**Figure S2.**
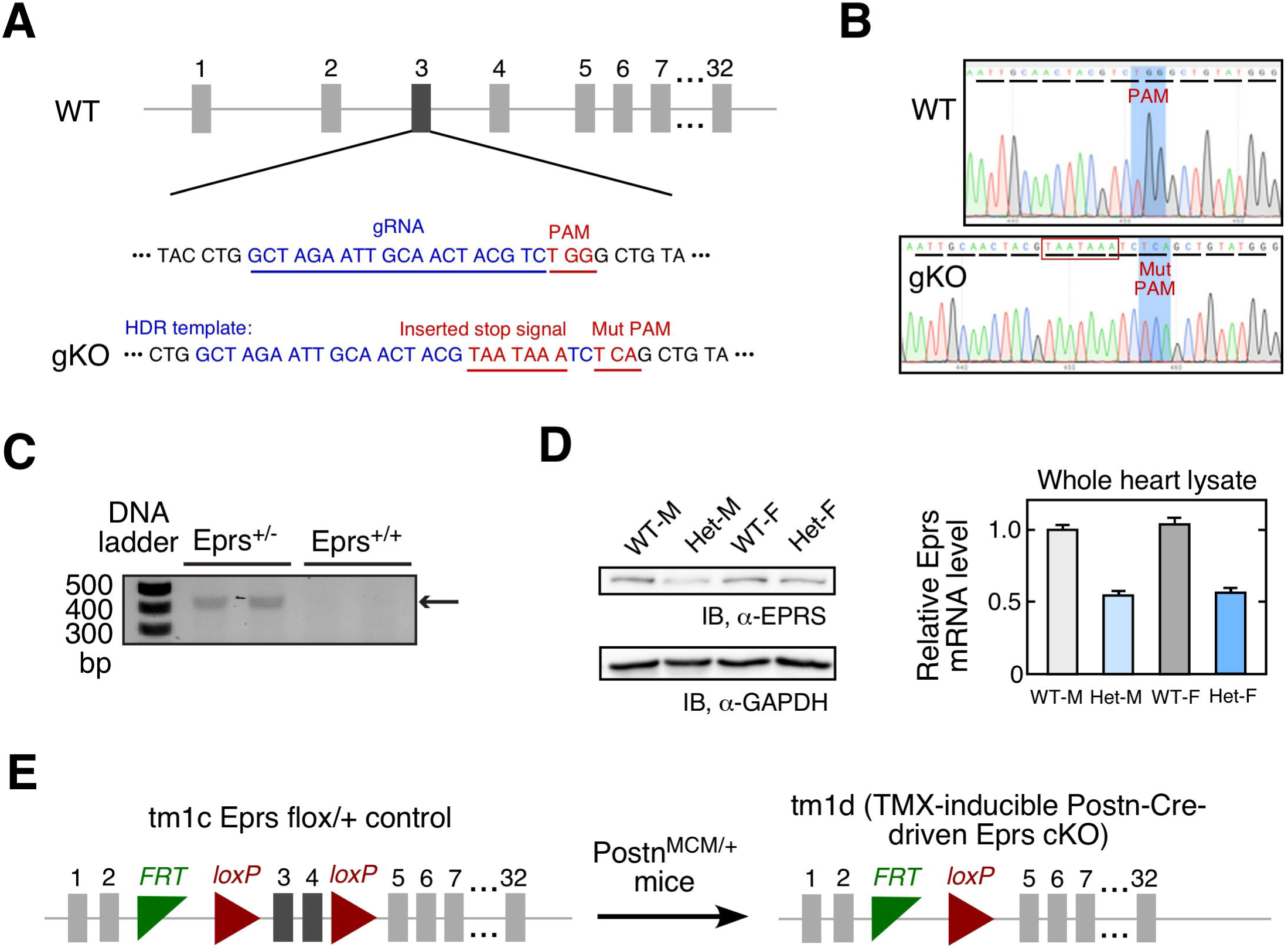
Generation and characterization of *Eprs* genetic knockout mice. **(A)** Schematic of the generation of *Eprs* global knockout mice by CRISPR technology. **(B)** Sanger DNA sequencing confirms an insertion of two tandem stop codons in *Eprs* knockout allele. **(C)** Genotyping of *Eprs*^+/+^ and *Eprs*^+/-^ mice. **(D)** *Eprs* mRNA and protein expression are reduced by ∼50% in *Eprs*^+/-^ mouse hearts. **(E)** Breeding strategy for generating tamoxifen-inducible Postn-Cre-driven *Eprs* conditional KO mice. ***: p≤0.001, NS: not significant by one-way ANOVA.

**Figure S3.**
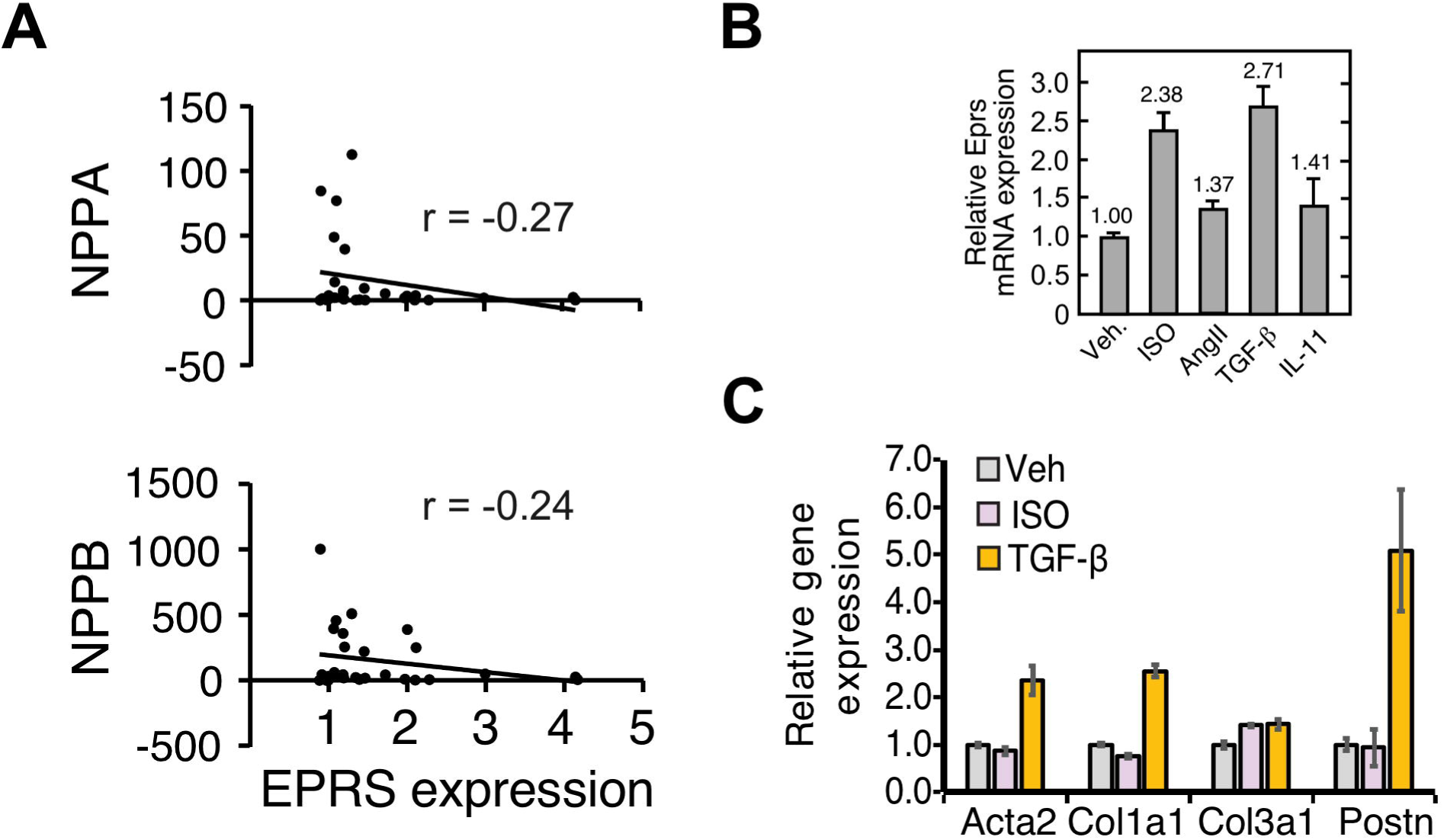
EPRS upregulation by multiple pro-hypertrophy and pro-fibrosis agonists in CF. **(A)** EPRS expression does not positively correlate with hypertrophic marker gene expression in human heart samples. **(B)** *Eprs* mRNA is induced by pro-hypertrophic and pro-fibrotic agonists in primary mouse adult CFs. **(C)** Myofibroblast marker gene mRNA expression is strongly induced by TGF-β, but not ISO in primary CFs.

**Figure S4.**
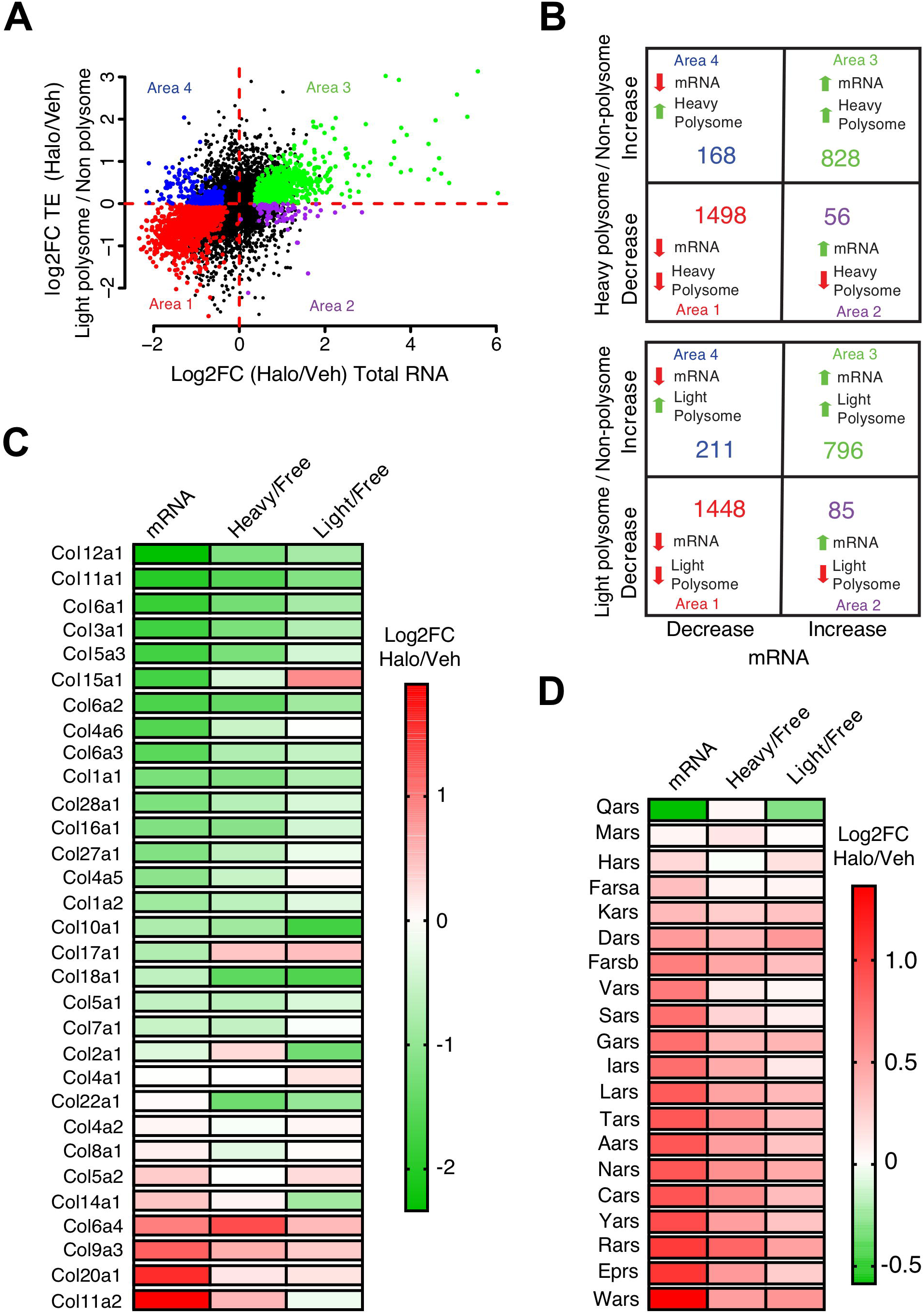
EPRS regulates Pro-rich protein expression in fibroblasts. **(A)** Differentially expressed genes identified by RNA-Seq and polysome-Seq are indicated by dot plot. Translation efficiency (TE) is indicated by the ratio between light polysome and polysome free fraction. The colored dots indicate statistically significantly changed genes in either RNA-Seq or polysome-Seq. The genes with p≤0.05 in either of three different groups (ratio of Halo vs. Veh. treated samples for total RNA, non polysome, and light polysome) were considered as significantly changed genes. All significantly changed genes were divided into four areas based on log2FC of total mRNAs and light polysome mRNAs. **(B)** Majority of genes show a synergistic change at mRNA and translational levels after EPRS inhibition by Halo. The number of genes is shown with changes at both translation efficiency (the ratio of heavy or light polysome to polysome free fraction) and steady-state mRNA levels in all 4 areas. **(C)** Log2FC of translation efficiency and steady-state mRNA level for all the collagens (typical Pro-rich genes as a positive control gene panel for Area 1). **(D)** Log2FC of translation efficiency and steady-state mRNA level for all cytosolic aminoacyl-tRNA synthetase (amino acid starvation markers as a positive control gene panel for Area 3).

**Figure S5.**
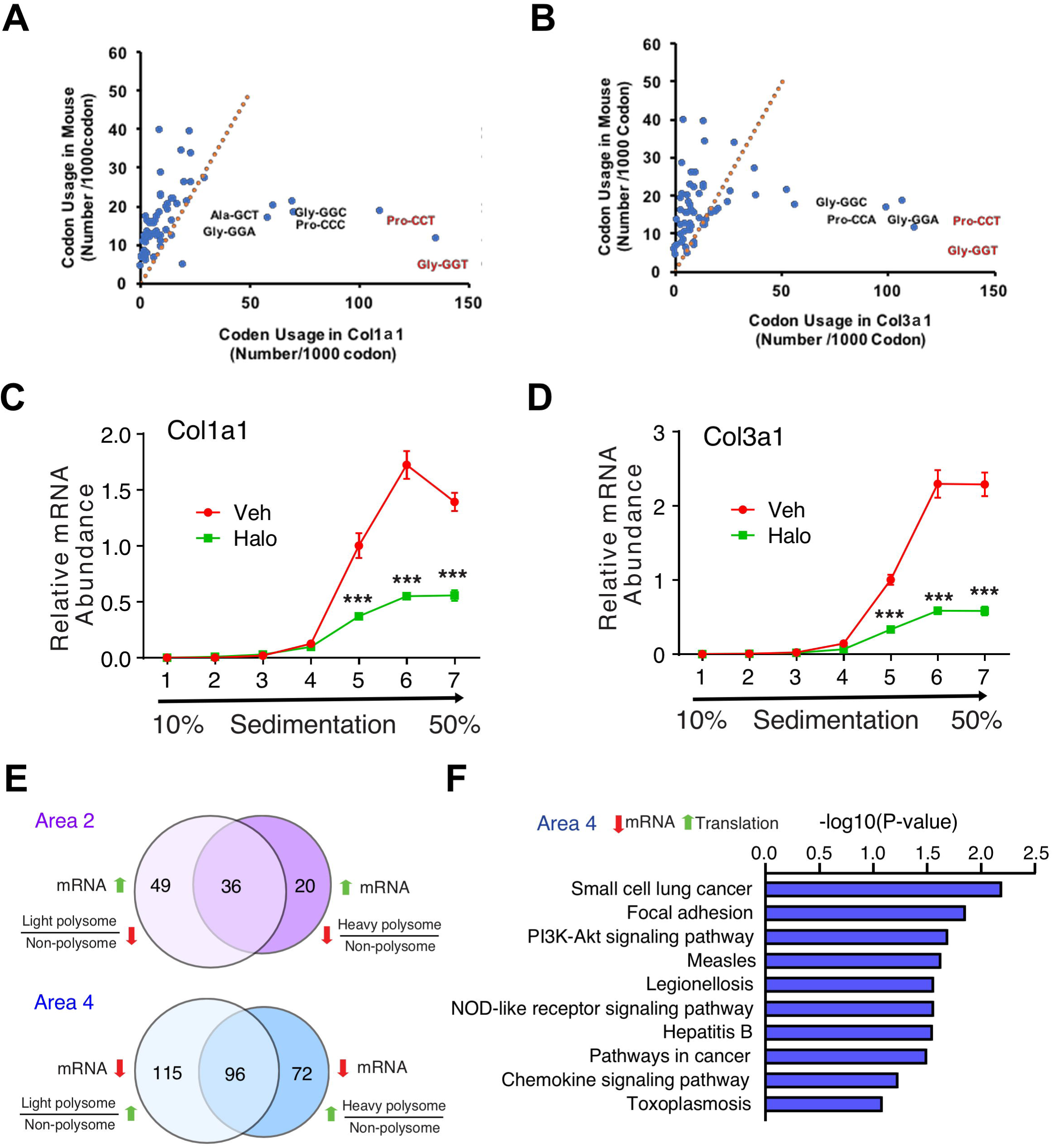
Halo downregulated collagen protein translation. **(A-B)** Genetic codon composition in mouse *Col1a1* and *Col3a1* genes and global mouse genome (https://www.genscript.com/tools/codon-frequency-table). **(C-D)** Halo reduced polysome association of collagen mRNAs in primary mouse CFs. **(E)** Translational dysregulated genes are indicated by overlapping changed genes in the same area of heavy polysome (Figure 4B) and light polysome **(Figure S4A)**. Genes decreased at the translational level and increased at steady-state mRNA level (Area 2) or increased at the translational level and decreased at steady-state mRNA level (Area 4) at translation and steady-state mRNA levels are shown. **(F)** KEGG pathway analyses of the gene cluster in area 4 from **(E)**.

**Figure S6.**
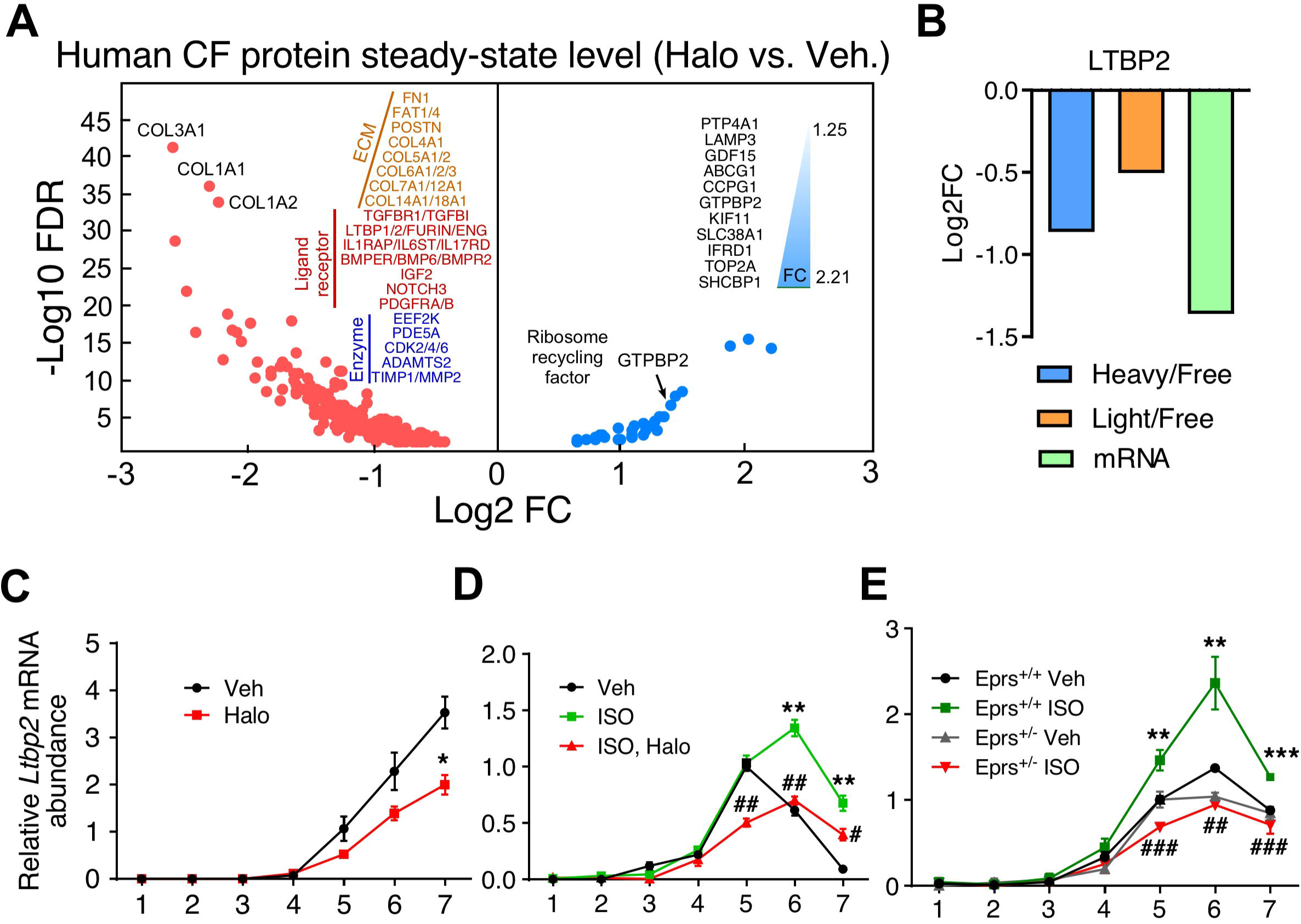
LTBP2 is a novel downstream target of EPRS. **(A)** Dot plot of mass spectrometry data of human CFs treated with 300 nM Halo from GSK company^15^. **(B)** *Ltbp2* mRNA level is reduced by Halo in both RNA-Seq and polysome-Seq. **(C)** Polysome associated *Ltbp2* mRNA level is reduced by Halo in primary CFs. **(D)** EPRS inhibition reverses ISO-induced polysome association of *Ltbp2* mRNA in CFs. *: Veh vs. ISO; #: ISO vs. ISO, Halo. **(E)** Polysome associated *Ltbp2* mRNA is reduced in *Eprs*^+/-^ CFs compared to WT (*Eprs*^+/+^) CFs after ISO treatment. *: *Eprs*^+/-^, Veh vs. ISO; #: *Eprs*^+/+^, ISO vs. Eprs^+/-^, ISO. *, #: p≤0.05, **, ##: p≤0.01, ***, ###: p≤0.001, NS: not significant by student t-test for two groups and one-way ANOVA for more than two groups.

**Figure S7.**
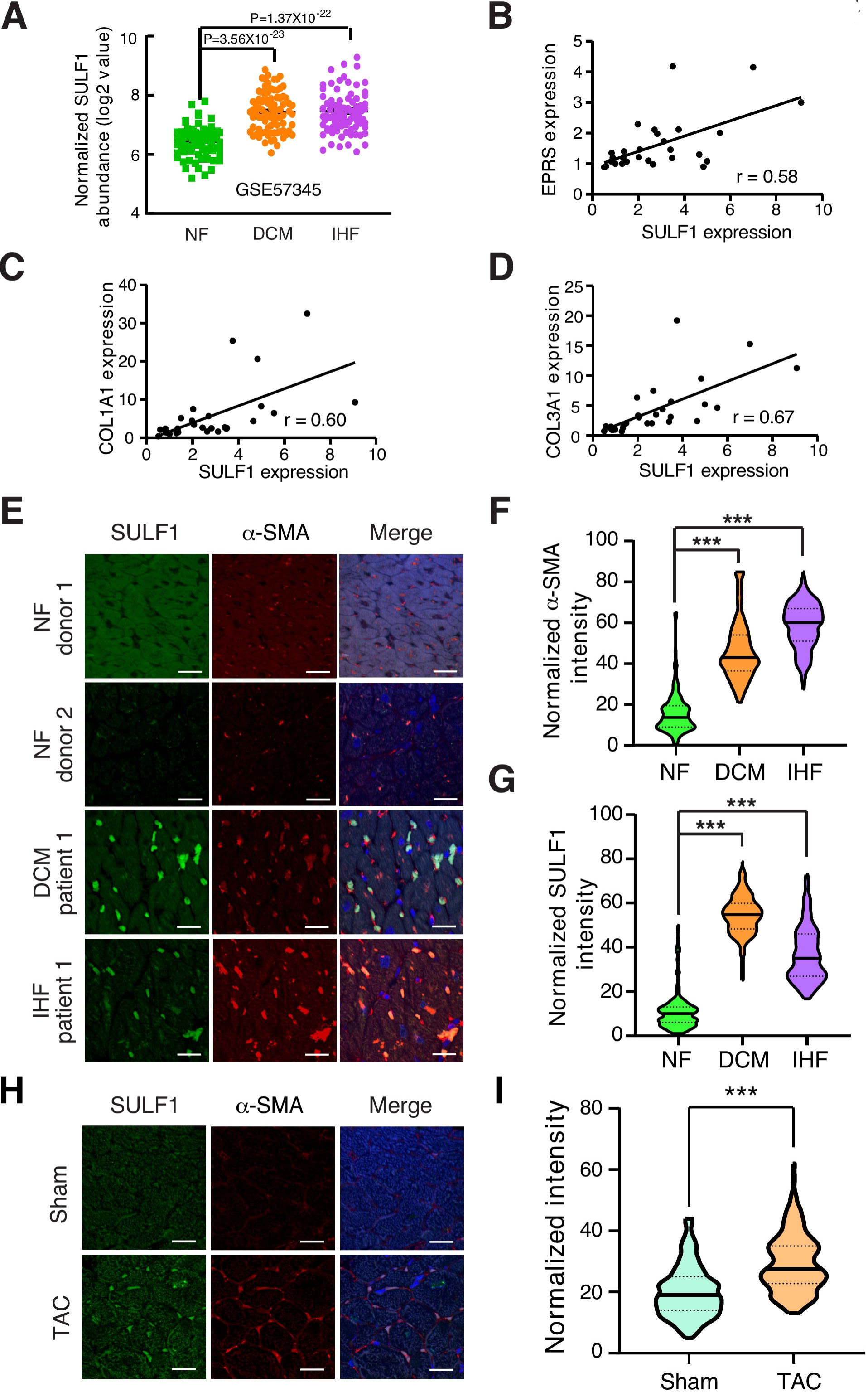
SULF1 is a human and mouse cardiac fibrosis marker. **(A)** *SULF1* mRNA expression is increased in dilated cardiomyopathy (DCM, n=82) and ischemic heart failure (IHF, n=95) patients compared to non-failing (NF, n=80) donor heart tissues from a public available microarray dataset GSE57345^35^. **(B-D)** The expression of SULF1 is correlated with EPRS **(B)**, COL1A1 **(C)**, and COL3A1 **(D)** in the validation cohort of human subjects. **(F)** Immunostaining shows SULF1 is highly expressed in α-SMA positive myofibroblasts in failing human hearts. Scale bar: 20 µm. **(F-G)** Quantitation of α-SMA **(F)** and SULF1 **(G)** protein expression in failing human and non-failing hearts in immunofluorescence staining in **(E)**. **(H-I)** Immunostaining shows SULF1 is highly expressed in α-SMA positive myofibroblasts in a TAC-induced mouse HF model **(H)** and quantitation of SULF1 intensity in **(I)**. Scale bar: 20 µm. **: p≤0.01, ***: p≤0.001 by unpaired two-tailed student t-test and one-way ANOVA in more than two groups.

**Figure S8.**
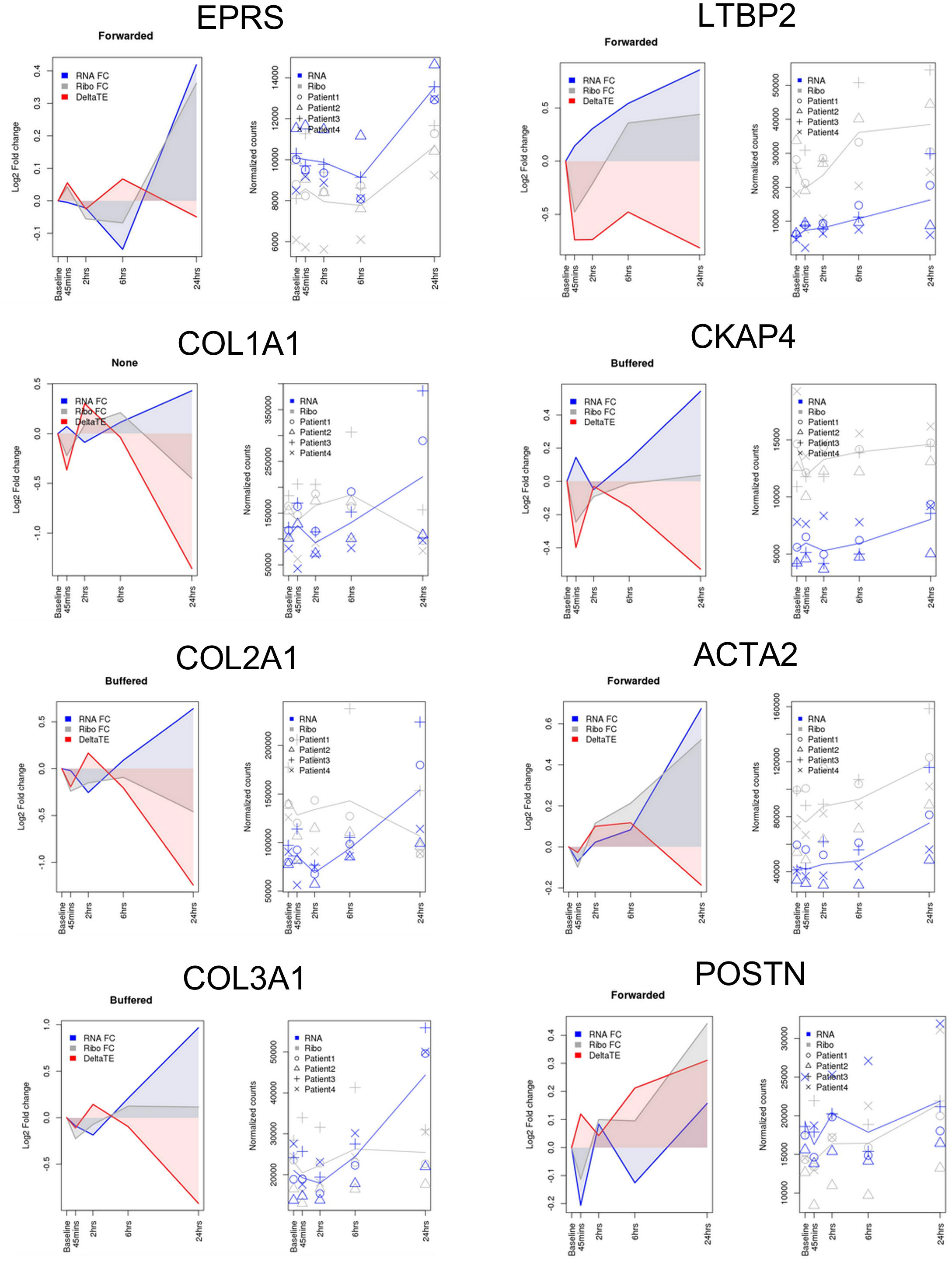
Fold change of gene expression at the levels of RNA-Seq (steady-state mRNA level), Ribo-Seq (ribosome footprints), and translation efficiency (DeltaTE= Ribo-Seq/RNA-Seq.

## ACKNOWLEDGMENTS

We are grateful to Omar Hedaya and Qiuqing Wang for critical reading of the manuscript and biostatistical consulting. We appreciate the technical assistance from Orazio Slivano (Aab CVRI) and Mohan Amy (Aab CVRI) in histology and surgical operations, respectively. NIH/3T3 cells were provided by Dr. Eric Small from Aab CVRI in URMC. Postn^MerCreMer^ mice were transferred from Eric Small lab and originally generated by Dr. Jeffery Molkentin from Cincinnati Children’s Hospital. Rommel Morales made efforts in detecting SULF1 protein level in human blood samples. *Eprs* global knockout mouse line was produced by Dr. Lin Gan from Mouse Genome Editing Core in URMC. None of the authors have any financial conflict of interest related to the research described in this manuscript.

## Materials and methods Reagents, antibodies and siRNAs

Halofuginone hydrochloride (Halo) was purchased from Santa Cruz Biotechnology (sc-211579). Isoprenaline (ISO) and cycloheximide (CHX) were purchased from Sigma-Aldrich. The siRNAs against mouse Sulf1 (assay ID: s109342 and s109343) and negative control siRNA were obtained from ThermoFisher Scientific and transfected into cardiac fibroblasts by lipofectamine 3000 (ThermoFisher) following the instruction manual.

Primary antibodies used in this study include: rabbit polyclonal anti-SULF1 antibody (PA5-50890) was purchased from ThermoFisher Scientific. Mouse monoclonal anti-COL1A1 (SAB1402151-100UG) and anti-a-smooth muscle actin (A2547) antibodies were purchased from Sigma-Aldrich. Mouse monoclonal anti-COL3A1 (sc-271249) was purchased from Santa Cruz Biotechnology. Mouse monoclonal anti-GAPDH (60004-1-Ig) was purchased from ProteinTech.

## Human specimens

All human samples of frozen cardiac tissues, including 17 samples from explanted failing hearts and 8 samples from non-failing donor hearts, as well as paraffin section slides from DCM, ISHF or non-failing hearts were acquired from the Cleveland Clinic. This study was approved by Material Transfer Agreement between the URMC and the Cleveland Clinic.

## Generation of *Eprs* global knockout mice

*Eprs* global knockout (gKO) mouse was generated by the Mouse Genome Editing Resource at URMC. The guide RNA (gRNA) used for Eprs KO was designed using a CRISPR RNA online design tool developed by Dr. Feng Zhang’s lab (http://crispr.mit.edu). The gRNA with highest score and lowest off-target probability was chosen. The efficiency of gRNA and Cas9-2xNLS (Synthego) were tested in a tube *in vitro* using a 500-bp mouse genomic DNA-derived PCR product that contains the target sequence. Then, 25 pmol of single guide RNA (sgRNA) and 25 pmol of Cas9 nuclease were mixed in the injection buffer in a total of 12.5 µl reaction. The mixture was then incubated at room temperature (RT) for 10 mins to form the RNP complex. Finally, this RNP complex was mixed with a single strand DNA template that contains two consecutive stop codons followed by an additional frame-shifting adenosine nucleotide (UAAUAAA) in frame of *Eprs* at 1:3 molar ratio for pronuclear injection. In this project, 148 injected embryos were transferred into 4 recipient C57/BL6J mice and finally we obtained 2 mosaic pups containing pre-mature stop codons in the exon 3 of *Eprs*. In order to get the heterozygous gKO mice for experiments, 2 mice from injection were bred with C57BL/6J WT mice for germline transmission and kept breeding with WT mice for 5 generations to remove the potential off-target sites. The genotyping primer set (Table S10) for the *Eprs* gKO mice was designed to complement with the mutated nucleotides of the protospacer adjacent motif (PAM) sequence. There is a 470-bp PCR amplification product from the mutant allele but no amplification from the WT allele.

Guide RNA: GCUAGAAUUGCAACUACGUCUGG

DNA template (157nt): GTTTACATAATGCTTCAATTCTAGG ACT GTG GCA TTC ACT GAC GTG AAT TCA ATC CTG CGC TAC CTG GCT AGA ATT GCA ACT ACG <u>TAA TAA A</u> TCT <u>CA</u>G CTG TAT GGG ACT AAC CTG ATG GAG CAC ACT GAGGTAAGCGAGGAGTATTTCTTTTCCT

## Generation of tamoxifen-inducible myofibroblast-specific *Eprs* conditional knockout mice

*Eprs* conditional knockout (cKO) mouse line *Eprs*_tm1c_B03 was purchased from The Center for Phenogenomics (TCP, Toronto, Canada) in the form of frozen sperms^1^. The *Eprs*^flox/+^ tm1c cKO mouse line was rederived using In Vitro Fertilization (IVF) was performed by the Mouse Genome Editing Resource at URMC. The *Eprs* cKO tm1c mouse line was bred with *Postn*^MCM/+^ mice^2^ to obtain tamoxifen-inducible myofibroblast-specific *Postn*-Cre-driven *Eprs*^flox/+^ tm1d cKO mouse line. We used one single initial dose of 30 µg/g of mouse body weight for tamoxifen (TMX) intraperitoneal injection followed by TMX food for the entire pathological stimulus of TAC surgery.

## Mouse heart failure models

In this study, we used two mouse heart failure models, including ISO osmotic minipump implantation and transverse aortic constriction (TAC) surgical model. Experimental mice are siblings generated from intercrosses between *Eprs*^+/-^ and C57BL/6J WT mice. Age and background matched WT and *Eprs*^+/-^ male mice at the age of ∼8-12 weeks were used for all the mouse studies. All the minipump implantation and surgeries were performed by the Mouse Microsurgical Core as previously described^3, 4^. At least six mice of each genetic background will be used for each treatment group to get statistically significant observations among different groups.

For the osmotic minipump implantation, mice were anesthetized using 2.0% isoflurane and placed on a heated surgical board. A side/upper back area skin incision was made, and the mini-osmotic pump was inserted subcutaneously and set to deliver ISO or vehicle at a rate of 20 mg/Kg/day. The incision was then closed with 6-0 coated vicryl in a subcuticular manner, and the animals were allowed to recover. The sutures were removed after 2 weeks since the pumps were transplanted. The pumps were not removed and remained for a time period of 4 weeks. The animals were euthanized after 4 weeks of ISO infusion and mouse hearts were harvested for experiments, including RNA, protein extraction and sectioning.

For the TAC surgery, mice were anesthetized via continuous administration of inhaled isofluorane (2%) while surgery was performed. The animals were placed supine, and a midline cervical incision was made to expose the trachea for direct intubation with 22-gauge plastic catheter. The catheter was connected to a volume-cycled ventilator supplying supportive oxygen. A right thoracotomy was performed. Stenosis was induced using a 27-gauge needle placed on the ascending aorta. Sham-operated mice underwent all aspects of the surgery besides the actual aortic ligature. A ligature was made around the needle and the aorta, completely occluding the aorta. The needle was then removed, causing severe aortic stenosis. Echocardiographic image collection was performed using a Vevo2100 echocardiography machine (VisualSonics, Toronto, Canada) and a linear-array 40 MHz transducer (MS-550D). Heart rate was monitored during echocardiography measurement. The stenotic gradient pressure was calculated to evaluate the efficacy of TAC. Left ventricular end-diastolic diameter (LVEDD), left ventricular end-systolic diameter (LVESD), wall thickness of left ventricular anterior (LVAWT) and posterior (LVPWT), ejection fraction (EF), and fractional shortening (FS) were assessed.

The mice are randomized for experiments. Animal operations, including ISO infusion, TAC surgery, and echocardiography measurement, were performed blindly by the Microsurgical Core surgeons. Sections and histology analysis were done by the Histology Core. For group size justification, we have performed a power analysis for One-way ANOVA. As an exemplary experiment, the standard deviation for weight after TAC or ISO treatment is about 10%. The minimum difference to be considered as significance is 25% in TAC- or ISO-induced cardiac hypertrophy and heart weight gain. With an overall type I error rate (alpha level) of 5%, at least 5 mice per treatment group are required to achieve 90% power to detect the difference of heart weight. In previous experiences from our Microsurgical Core, we have observed a survival rate of ∼90% after the TAC procedure. To offset the possible loss of one mouse per treatment, we used at least 6 mice per treatment group.

## Adult cardiomyocyte and cardiac fibroblasts isolation

Langendorff perfusion system was used to isolate adult cardiomyocytes (CMs) and cardiac fibroblasts from the murine heart. Mice were fully anesthetized via intraperitoneal injection of ketamine/xylazine. Once losing pedal reflex, the mouse was secured in a supine position. The heart was excised and fastened onto the CM perfusion apparatus and perfusion was initiated in the Langendorff mode. Our Langendorff perfusion and digestion consisted of three steps at 37°C: 4 mins with perfusion buffer (0.6 mM KH_2_PO_4_, 0.6 mM Na_2_HPO_4_, 10 mM HEPES, 14.7 mM KCl, 1.2 mM MgSO_4_, 120.3 mM NaCl, 4.6 mM NaHCO_3_, 30 mM taurine, 5.5 mM glucose, and 10 mM 2,3-butanedione monoxime), then switched to digestion buffer (300 U/ml collagenase II [Worthington] in perfusion buffer) for 3 mins, and finally perfused with digestion buffer supplemented with 40 µM CaCl_2_ for 8 mins. After perfusion, the ventricle was placed in sterile 35 mm dish with 2.5 ml digestion buffer and shredded into several pieces with forceps. 5 ml stopping buffer (10% FBS, 12.5 µM CaCl_2_ in perfusion buffer) was added and pipetted several times until tissues disperse readily, and solution turned cloudy. The cell solution was passed through 100 µm strainer. CMs were settled by incubating the cell suspension at 37°C for 30 mins. The CMs were resuspended in 10 ml stopping buffer and subjected to several steps of calcium ramping: 100 µM CaCl_2_, 2 mins; 500 µM CaCl_2_, 4 mins; 1.4 mM CaCl_2_, 7 mins. Then the CMs were seeded onto a glass bottom dish (Nest Biotechnology) pre-coated with laminin (ThermoFisher Scientific). Plates were centrifuged for 5 mins at 1,000 g at 4°C to increase the adherence, cultured at 37°C for ∼1 hr, and then switched to CM media (MEM [Corning] with 0.2% BSA, 10 mM HEPES, 4 mM NaHCO3, 10 mM creatine monohydrate, 1% penicillin/streptomycin, 0.5% insulin-selenium-transferrin and blebbistatin) for cell culture and further treatments.

Cardiac fibroblasts (CFs) from the supernatant are pelleted for 5 mins at 1,000 rpm at 4°C. CFs were plated in 4-5 ml CF media (DMEM with 10% FBS and 1% penicillin/streptomycin) in 60 mm plate and washed vigorously 3-5 times with 2 ml 1x PBS after 2-3 hrs, and replaced with fresh CF media. For CF only isolation, pre-weened mice were fully anesthetized and the heart was directly cut into small pieces and digested in the digestion buffer for 4x 10 mins at 37°C with slow stirring and CFs were plated the same as Langendorff isolation of CFs.

## Cell culture

NIH/3T3 cells were a gift from Eric Small lab and cultured in DMEM supplemented with 10% bovine calf serum (VWR) and 1% penicillin/streptomycin (ThermoFisher). For halofuginone treatment, NIH/3T3 cells were grown to ∼60-70% confluency and treated with 100 nM halofuginone or vehicle, or at the concentration as indicated for 24 hrs. Primary CFs isolated from mouse hearts were cultured in DMEM supplemented with 10% FBS (ThermoFisher) and 1% penicillin/streptomycin. Primary cells were used before P2 generation for polysome profiling and at P0 for CF activation assays. siRNA transfection (100 nM) in primary CFs was performed using lipofectamine 3000 following the manufacturer’s instructions.

## Polysome profiling

Polysome profiling was performed to measure the global cytosolic translational status or the translational efficiency of specific genes in both NIH/3T3 cells and primary CFs as previously described^5^. Briefly, after drug treatment of cells (Halo, ISO or Veh), cycloheximide (CHX, Sigma) was added to the cells at a final concentration of 100 µg/ml for 15 mins before lysis to freeze ribosomes on mRNAs. ∼5 × 10^6^ cells were lysed in TMK lysis buffer (10 mM Tris-HCl pH 7.4, 5 mM MgCl_2_, 100 mM KCl, 1% Triton X-100, 0.5% Deoxycholate, 2 mM DTT) containing 100 µg/ml CHX, 4 U/ml RNase inhibitor (NEB), and proteinase inhibitor cocktail (Roche) on ice for 15 mins. Equal amounts of A_260_ absorbance from each sample were loaded onto a 10-50% sucrose gradient solution, and centrifuged at 29,000 rpm for 4 hrs at 4°C. The polysome fractions were collected from each sample by a density gradient fractionation system (BRANDEL). For specific genes, total RNA was extracted from the same volume of each fraction with Trizol LS (ThermoFisher) and used for RT-qPCR. Renilla luciferase mRNA made by *in vitro* transcription was used as a spike-in loading control. Based on the UV absorbance curve, the 22 fractions were pooled into 7 samples for RT-qPCR analysis including free mRNP, 40S small ribosome subunit, 60S large ribosome subunit, 80S monosome, light polysomes (di-ribosome, tri-ribosome, etc.), and heavy polysomes (>5 ribosomes). For polysome-Seq, all the fractions were pooled into 3 samples, polysome free (free mRNP, 40S, 60S subunit), light polysomes (monosome, disome, trisome, etc.), and heavy polysomes (>5 ribosomes). Total RNAs were extracted from the same volume of 3 samples and subjected to RNA-Seq. The RNA-Seq and polysome-Seq data were uploaded to NCBI GEO database with ID of GSE136838.

## mRNA expression

For heart tissues (human and mouse) or cell samples, the RNA extraction was performed using TRIzol reagent (ThermoFisher) following instructions in the manual and used for the detection of the expression of specific genes. Briefly, the tissues were homogenized in TRIzol using Minilys Personal Homogenizer (Bertin Technologies) and placed on ice for 15 mins to lyse the tissue. For the polysome fraction samples, total RNA extraction was performed using TRIzol LS reagent at a volume ratio of 1:3 following the manual. The *in vitro* transcribed renilla luciferase mRNA was added to the samples before RNA extraction as a spike-in loading control.

For the mRNA detection, 1 µg of total RNA was used as a template for reverse transcription using the iScript cDNA Synthesis Kit (Bio-Rad). cDNA was used for detecting the expression of Eprs, Sulf1 and the marker genes, Nppa, Nppb, Col1a1 and Col3a1. 18S rRNA or Gapdh was used as a normalization control for mRNA expression. The SYBR Green primer sequences or the Taqman probes were listed in Table S10.

## RNA-Seq and polysome-Seq NGS data processing and alignment

Total RNA extracted from halofuginone or vehicle treated NIH/3T3 cells or polysome fractions were treated with DNase I (NEB) to remove potential genomic DNA in the RNA samples. The DNase I treated RNA samples were purified with phenol:chloroform:isoamyl alcohol and then subjected to RNA-Seq at the Genomic Research Center of URMC. Raw reads generated from the Illumina HiSeq2500 sequencer were demultiplexed using bcl2fastq version 2.19.0. Quality filtering and adapter removal are performed using Trimmomatic version 0.36^6^ with the following parameters: “TRAILING:13 LEADING:13 ILLUMINACLIP: adapters.fasta:2:30:10 SLIDINGWINDOW:4:20 MINLEN:15”. Processed/cleaned reads were then mapped to the *Mus musculus* reference genome (GRCm38, mg38) with STAR_2.5.2b^7^ given the following parameters: “—twopassMode Basic --runMode alignReads --genomeDir ${GENOME} -- readFilesIn ${SAMPLE} --outSAMtype BAM SortedByCoordinate --outSAMstrandField intronMotif --outFilterIntronMotifs RemoveNoncanonical”. The subread-1.5.0^8^ package (featureCounts) was used to derive gene counts given the following parameters: “-s 2 -t exon -g gene_name” and the gencode M12 gene annotations. Differential expression analysis and data normalization was performed using DESeq2-1.16.1^9^ with an adjusted p-value threshold of 0.05 within an R-3.4.1^10^ environment. A batch factor was given to the differential expression model in order to control for batch differences. Gene ontology and KEGG pathway enrichment analyses were performed using DAVID Bioinformatics Resources 6.8^11^.

## Proline motif analysis across mouse and human proteomes

The proteomic sequences of UniProtKB/Swiss-Prot reviewed proteins of Homo sapiens (UP000005640, reviewed, 20,328 sequences) and Mus musculus (UP000000589, reviewed, 16,966 sequences) were downloaded from Uniprot database (https://www.uniprot.org/) in fasta format. Each protein sequence was processed and the number of consecutive prolines (PP, PPP, PPPP, P…P_n_ [n=2-9, and n≥10], etc.) motifs were calculated using R package “Biostrings” (R version 3.3.3). The number of consecutive proline motifs were quantified as previous described^12^. Briefly, the consecutive proline motifs were counted without any overlapping and the number of PP or PPP within protein containing higher number of consecutive proline (more than 2 or 3 for PP or PPP) was determined by the whole number after division of the consecutive proline length by 2 (PP) or 3 (PPP). For example, the 4 and 5 consecutive proline stretch will be considered as 2 PP motifs and 1 PPP motifs while 6 and 7 consecutive proline stretch as 3 PP motifs and 2 PPP motifs. The conserved PP motif containing genes were generated by overlapping the PP motif containing genes in human and mouse proteomes.

## Cardiac fibroblasts activation assay

Adult CF cells were isolated from hearts of pre-weened mice and seeded into 35 mm glass bottom dishes. After attachment for 2 hrs, the cells were washed with PBS for 3 times, replaced with fresh CF growth media (DMEM supplemented with 10% FBS and 1% penicillin/streptomycin), and cultured at 37°C overnight. On the second day, 100 nM of Sulf1 siRNAs or control siRNAs were transfected using lipofectamine 3000 following the manual. The cells were cultured for additional 36 hrs and 10 ng/ml recombinant mouse TGFβ1 (ThermoFisher) was added to the media for 24 hrs following a 12 hr serum starvation. Then the cells were fixed with 4% paraformaldehyde (PFA) for immunofluorescence or lysed in TRIzol reagent for the marker gene detection by RT-qPCR.

## Immunofluorescence and immunohistochemistry

For the immunofluorescence, the primary CF cells were fixed using 4% PFA/PBS and washed with PBS for 3x 5 min. The cells were permeabilized by ice-cold 0.5% triton X-100/PBS for 5 mins and washed with PBS for 3x 5 min. After blocking with 4% BSA in PBS for 30 mins, the cells were incubated with indicated primary antibodies (rabbit anti-SULF1, 1:200; mouse anti-α-SMA, 1:500; mouse anti-COL1A1, 1:200) in blocking solution (4% BSA in PBS) overnight at 4°C, and washed with PBS for 3x 5 min. Then, the cells were co-stained with the Alex Fluor-488 and Alex Fluor-594 conjugated secondary antibodies (ThermoFisher, 1:250) in blocking solution for 45 mins and washed with PBS for 3x 5 min. Finally, the cells were co-stained with DAPI for 5 mins and kept in PBS buffer inside before imaging.

For the immunohistochemistry, the mouse heart tissues were fixed with 10% neutralized formalin solution and processed for paraffin embedded sections in the Histological Core of Aab CVRI. For paraffin embedded sections from both human and mouse hearts, the sections were deparaffinized by the following steps: xylene (100%) for 2x 5 min; ethanol (100%) for 2x 5 min; ethanol (95%) for 1x 5 min; ddH_2_O for 2x 5 min. Antigen retrieval was performed by boiling the deparaffinized section in 10 mM citrate buffer, pH 6.0 and auto-fluorescence was quenched by incubation in 3% H_2_O_2_/PBS for 30 mins at RT. Then the slides were incubated with blocking buffer (5% goat serum in PBS) for 1 hr at RT and the indicated primary antibody (rabbit anti-SULF1, 1:200; mouse anti-a-SMA, 1:500; mouse anti-COL1A1, 1:200) in blocking solution overnight at 4°C. Alex Fluor-488 and Alex Fluor-594 conjugated secondary antibodies were incubated for 1 hr following 3x 5 min washing step. Finally, the slides were co-stained with DAPI in the mounting media and covered by coverslips. The images were taken using an Olympus FV1000 confocal microscope and the intensity was measured by NIH Image J software.

## Wheat germ agglutinin (WGA) and phalloidin staining

WGA staining was used to quantify the size of CMs in the murine heart after drug treatments. Deparaffinization, antigen retrieval and quenching of auto-fluorescence were performed the same as described above. The section dots were marked by Dako pen and stained with 10 µg/ml WGA-Alex Fluor-488 (ThermoFisher) for 1.5 hr at RT. Then the slides were washed with PBS for 3x 5 min. The slides were covered by coverslips with antifade solution (containing DAPI) for imaging. The images were taken in the fluorescence microscope and Image J software was used to quantify the cell size of CMs.

For isolated primary mouse CMs, cultured myocytes were fixed using 4% paraformaldehyde, washed with PBS, and permeabilized in 0.5% Triton X-100. Alexa Fluor^TM^ 594 Phalloidin (ThermoFisher, Cat. No. A12381) was used to stain the skeleton actin protein for 30 min at RT. The stained cells were gently washed with PBS for 3x 5 min and mounted using anti-fade mounting medium with DAPI (Vector). The images were taken in the fluorescence microscope and Image J software was used to quantify the cell size of CMs.

## Picrosirius red staining

Picrosirius staining was performed to measure the cardiac fibrosis in heart failure models using picrosirius red solution (Abcam) following the manufacturer’s instruction. Briefly, paraffin embedded tissue sections were deparaffinized and incubated in picrosirius red solution at RT for 1 hr. Then, slides were subjected to 2 washes of 1% acetic acid and 100% of ethyl alcohol, and mounted in a mounting medium. Images were captured using the PrimeHisto XE Histology Slide Scanner (Carolina) and cardiac fibrotic area was quantified from the whole heart images of picrosirius red staining using Image J software.

## Statistical Analysis

All quantitative data were presented as mean ± SEM, and analyzed using Prism 8 software (GraphPad). For comparison between 2 groups, a Student t-test was performed. For multiple comparisons among ≥3 groups, 1-way ANOVA was performed. Statistical significance was assumed at a value of P<0.05.

---

**Table S2:**
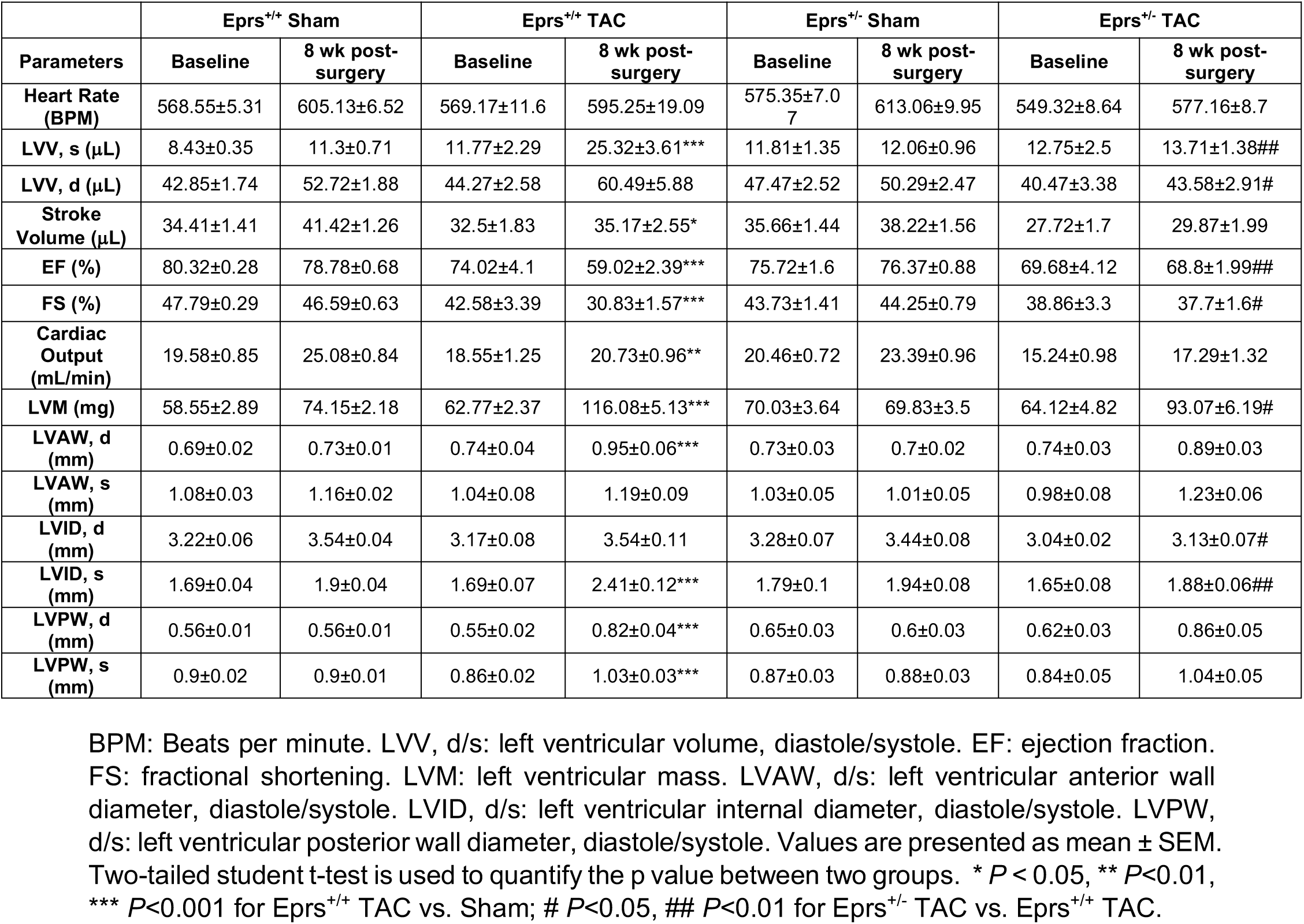
Summary of echocardiograph data in Eprs^+/+^ and Eprs^+/-^ mice after 8 weeks of Sham or TAC surgeries.

**Table S10.**
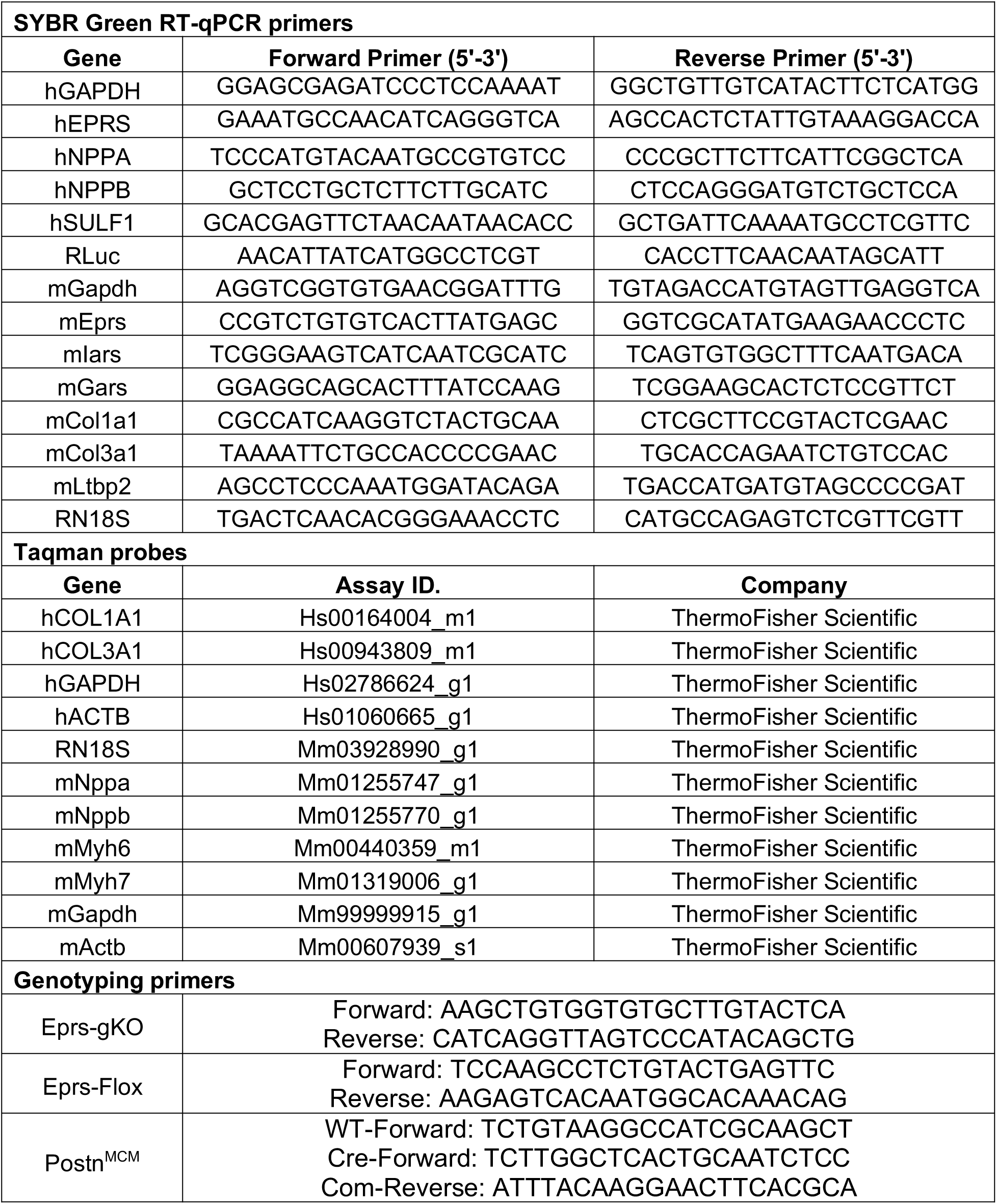
SYBR Green primers and Taqman probes used for RT-qPCR in this research.

**Table S1:** Reanalysis of mRNA expression of all the translation machinery component genes in TGF-β-versus vehicle-treated human cardiac fibroblasts from published database (*Nature* 2017 552(7683): 110-115).

**Table S3:** Log2FC of mRNAs in RNA-Seq and 3 different pooled fractions of polysome-Seq upon halofuginone treatment. RNA-Seq and polysome-Seq were performed in 100 nM Halo or Veh. treated fibroblast cells. The log2FC and p value were calculated for all the mRNAs between Halo and Veh. in all four groups (total RNA, non polysome, light polysome, and heavy polysome).

**Table S4:** Differentially expressed genes in four areas of gene clusters. The genes with p≤0.05 in either of four different groups (ratio of Halo vs. Veh. treated samples for total RNA, ribosome free, light polysome, and heavy polysome) were considered as significantly changed genes. All significantly changed genes were divided into four areas based on log2FC of total mRNAs and polysome mRNAs (heavy or light polysome). Area 1: total mRNA decreased and polysomal mRNA decreased; Area 2: total mRNA increased and polysomal mRNA decreased; Area 3: total mRNA increased and polysomal mRNA increased; Area 4: total mRNA increased and polysomal mRNA decreased.

**Table S5:** KEGG pathway analyses in four different areas of gene clusters. All the genes in four areas were subjected to KEGG pathway analyses in DAVID Bioinformatics Resources and the significantly enriched genes were listed. Note: No KEGG pathways was significantly enriched in Area 4 due to a limited number of genes in this area.

**Table S6:** Proline motifs analyses in area 1 and 3. The number of proline motifs were quantified in the genes of area 1 and 3.

**Table S7:** The list of PP motif containing genes that were quantified as conserved hits across human and mouse proteomes. The proline motifs were quantified across mouse and human proteomes and the genes containing PP motif in both human and mouse were considered as conserved PP motif containing genes.

**Table S8:** The list of genes containing PP motifs and undergoing downregulation at polysomal mRNA and steady state protein levels. By overlapping of human/mouse conserved PP motif containing genes, Halo-triggered polysomal mRNA-downregulated genes, and mass spectrometry-detected protein-decreased genes, 83 common hits were significantly downregulated by low dose of Halo at the translational level.

**Table S9:** GO analysis of the overlapped 83 genes. Gene ontology analysis was performed for the 83 overlapped genes and fold change of all genes in top 5 GO_BP terms were listed.

## Supplemental sequence information

**Human SULF1 protein sequence:** MKYSCCALVLAVLGTELLGSLCSTVRSPRFRGRIQQERKNIRPNIILVLTDDQDVELGSLQVMNKTR KIMEHGGATFINAFVTTPMCCPSRSSMLTGKYVHNHNVYTNNENCSSPSWQAMHEPRTFAVYLN NTGYRTAFFGKYLNEYNGSYIPPGWREWLGLIKNSRFYNYTVCRNGIKEKHGFDYAKDYFTDLITN ESINYFKMSKRMYPHRPVMMVISHAAPHGPEDSAPQFSKLYPNASQHITPSYNYAPNMDKHWIM QYTGPMLPIHMEFTNILQRKRLQTLMSVDDSVERLYNMLVETGELENTYIIYTADHGYHIGQFGLVK GKSMPYDFDIRVPFFIRGPSVEPGSIVPQIVLNIDLAPTILDIAGLDTPPDVDGKSVLKLLDPEKPGNR FRTNKKAKIWRDTFLVERGKFLRKKEESSKNIQQSNHLPKYERVKELCQQARYQTACEQPGQKW QCIEDTSGKLRIHKCKGPSDLLTVRQSTRNLYARGFHDKDKECSCRESGYRASRSQRKSQRQFL RNQGTPKYKPRFVHTRQTRSLSVEFEGEIYDINLEEEEELQVLQPRNIAKRHDEGHKGPRDLQASS GGNRGRMLADSSNAVGPPTTVRVTHKCFILPNDSIHCERELYQSARAWKDHKAYIDKEIEALQDKI KNLREVRGHLKRRKPEECSCSKQSYYNKEKGVKKQEKLKSHLHPFKEAAQEVDSKLQLFKENNR RRKKERKEKRRQRKGEECSLPGLTCFTHDNNHWQTAPFWNLGSFCACTSSNNNTYWCLRTVNE THNFLFCEFATGFLEYFDMNTDPYQLTNTVHTVERGILNQLHVQLMELRSCQGYKQCNPRPKNLD VGNKDGGSYDLHRGQLWDGWEG

**Mouse SULF1 protein sequence:** MKYSLWALLLAVLGTQLLGSLCSTVRSQRFRGRIQQERKNIRPNIILVLTDDQDVELGSLQVMNKT RKIMEQGGATFTNAFVTTPMCCPSRSSMLTGKYVHNHNVYTNNENCSSPSWQAMHEPRTFAVY LNNTGYRTAFFGKYLNEYNGSYIPPGWREWLGLIKNSRFYNYTVCRNGIKEKHGFDYAKDYFTDLI TNESINYFKMSKRMYPHRPIMMVISHAAPHGPEDSAPQFSKLYPNASQHITPSYNYAPNMDKHWI MQYTGPMLPIHMEFTNVLQRKRLQTLMSVDDSVERLYNMLVESGELDNTYIIYTADHGYHIGQFG LVKGKSMPYDFDIRVPFFIRGPSIEPGSIVPQIVLNIDLAPTILDIAGLDSPSDVDGKSVLKLLDLEKPG NRFRTNKKAKIWRDTFLVERGKFLRKKEESGKNIQQSNHLPKYERVKELCQQARYQTACEQPGQ NWQCIEDTSGKLRIHKCKGPSDLLTVRQNARNLYSRGLHDKDKECHCRDSGYRSSRSQRKNQR QFLRNKGTPKYKPRFVHTRQTRSLSVEFEGEIYDINLEEEELQVLPPRSIAKRHDEGHQGFIGHQAA AGDIRNEMLADSNNAVGLPATVRVTHKCFILPNDTIHCERELYQSARAWKDHKAYIDKEIEVLQDKI KNLREVRGHLKKRKPEECGCGDQSYYNKEKGVKRQEKLKSHLHPFKEAAAQEVDSKLQLFKEHR RRKKERKEKKRQRKGEECSLPGLTCFTHDNNHWQTAPFWNLGSFCACTSSNNNTYWCLRTVNE THNFLFCEFATGFLEYFDMNTDPYQLTNTVHTVERSILNQLHIQLMELRSCQGYKQCNPRPKSLDI GAKEGGNYDPHRGQLWDGWEG

**Human COL1A1 protein sequence:** MFSFVDLRLLLLLAATALLTHGQEEGQVEGQDEDIPPITCVQNGLRYHDRDVWKPEPCRICVCDN GKVLCDDVICDETKNCPGAEVPEGECCPVCPDGSESPTDQETTGVEGPKGDTGPRGPRGPAGPP GRDGIPGQPGLPGPPGPPGPPGPPGLGGNFAPQLSYGYDEKSTGGISVPGPMGPSGPRGLPGP PGAPGPQGFQGPPGEPGEPGASGPMGPRGPPGPPGKNGDDGEAGKPGRPGERGPPGPQGAR GLPGTAGLPGMKGHRGFSGLDGAKGDAGPAGPKGEPGSPGENGAPGQMGPRGLPGERGRPG APGPAGARGNDGATGAAGPPGPTGPAGPPGFPGAVGAKGEAGPQGPRGSEGPQGVRGEPGPP GPAGAAGPAGNPGADGQPGAKGANGAPGIAGAPGFPGARGPSGPQGPGGPPGPKGNSGEPG APGSKGDTGAKGEPGPVGVQGPPGPAGEEGKRGARGEPGPTGLPGPPGERGGPGSRGFPGAD GVAGPKGPAGERGSPGPAGPKGSPGEAGRPGEAGLPGAKGLTGSPGSPGPDGKTGPPGPAGQ DGRPGPPGPPGARGQAGVMGFPGPKGAAGEPGKAGERGVPGPPGAVGPAGKDGEAGAQGPP GPAGPAGERGEQGPAGSPGFQGLPGPAGPPGEAGKPGEQGVPGDLGAPGPSGARGERGFPGE RGVQGPPGPAGPRGANGAPGNDGAKGDAGAPGAPGSQGAPGLQGMPGERGAAGLPGPKGD RGDAGPKGADGSPGKDGVRGLTGPIGPPGPAGAPGDKGESGPSGPAGPTGARGAPGDRGEPG PPGPAGFAGPPGADGQPGAKGEPGDAGAKGDAGPPGPAGPAGPPGPIGNVGAPGAKGARGSA GPPGATGFPGAAGRVGPPGPSGNAGPPGPPGPAGKEGGKGPRGETGPAGRPGEVGPPGPPGP AGEKGSPGADGPAGAPGTPGPQGIAGQRGVVGLPGQRGERGFPGLPGPSGEPGKQGPSGASG ERGPPGPMGPPGLAGPPGESGREGAPGAEGSPGRDGSPGAKGDRGETGPAGPPGAPGAPGA PGPVGPAGKSGDRGETGPAGPAGPVGPVGARGPAGPQGPRGDKGETGEQGDRGIKGHRGFSG LQGPPGPPGSPGEQGPSGASGPAGPRGPPGSAGAPGKDGLNGLPGPIGPPGPRGRTGDAGPV GPPGPPGPPGPPGPPSAGFDFSFLPQPPQEKAHDGGRYYRADDANVVRDRDLEVDTTLKSLSQ QIENIRSPEGSRKNPARTCRDLKMCHSDWKSGEYWIDPNQGCNLDAIKVFCNMETGETCVYPTQ PSVAQKNWYISKNPKDKRHVWFGESMTDGFQFEYGGQGSDPADVAIQLTFLRLMSTEASQNITY HCKNSVAYMDQQTGNLKKALLLQGSNEIEIRAEGNSRFTYSVTVDGCTSHTGAWGKTVIEYKTTK TSRLPIIDVAPLDVGAPDQEFGFDVGPVCFL

**Mouse COL1A1 protein sequence:** MFSFVDLRLLLLLGATALLTHGQEDIPEVSCIHNGLRVPNGETWKPEVCLICICHNGTAVCDDVQC NEELDCPNPQRREGECCAFCPEEYVSPNSEDVGVEGPKGDPGPQGPRGPVGPPGRDGIPGQPG LPGPPGPPGPPGPPGLGGNFASQMSYGYDEKSAGVSVPGPMGPSGPRGLPGPPGAPGPQGFQ GPPGEPGEPGGSGPMGPRGPPGPPGKNGDDGEAGKPGRPGERGPPGPQGARGLPGTAGLPG MKGHRGFSGLDGAKGDAGPAGPKGEPGSPGENGAPGQMGPRGLPGERGRPGPPGTAGARGN DGAVGAAGPPGPTGPTGPPGFPGAVGAKGEAGPQGARGSEGPQGVRGEPGPPGPAGAAGPAG NPGADGQPGAKGANGAPGIAGAPGFPGARGPSGPQGPSGPPGPKGNSGEPGAPGNKGDTGA KGEPGATGVQGPPGPAGEEGKRGARGEPGPSGLPGPPGERGGPGSRGFPGADGVAGPKGPSG ERGAPGPAGPKGSPGEAGRPGEAGLPGAKGLTGSPGSPGPDGKTGPPGPAGQDGRPGPAGPP GARGQAGVMGFPGPKGTAGEPGKAGERGLPGPPGAVGPAGKDGEAGAQGAPGPAGPAGERG EQGPAGSPGFQGLPGPAGPPGEAGKPGEQGVPGDLGAPGPSGARGERGFPGERGVQGPPGPA GPRGNNGAPGNDGAKGDTGAPGAPGSQGAPGLQGMPGERGAAGLPGPKGDRGDAGPKGAD GSPGKDGARGLTGPIGPPGPAGAPGDKGEAGPSGPPGPTGARGAPGDRGEAGPPGPAGFAGP PGADGQPGAKGEPGDTGVKGDAGPPGPAGPAGPPGPIGNVGAPGPKGPRGAAGPPGATGFPG AAGRVGPPGPSGNAGPPGPPGPVGKEGGKGPRGETGPAGRPGEVGPPGPPGPAGEKGSPGAD GPAGSPGTPGPQGIAGQRGVVGLPGQRGERGFPGLPGPSGEPGKQGPSGSSGERGPPGPMGP PGLAGPPGESGREGSPGAEGSPGRDGAPGAKGDRGETGPAGPPGAPGAPGAPGPVGPAGKNG DRGETGPAGPAGPIGPAGARGPAGPQGPRGDKGETGEQGDRGIKGHRGFSGLQGPPGSPGSP GEQGPSGASGPAGPRGPPGSAGSPGKDGLNGLPGPIGPPGPRGRTGDSGPAGPPGPPGPPGP PGPPSGGYDFSFLPQPPQEKSQDGGRYYRADDANVVRDRDLEVDTTLKSLSQQIENIRSPEGSRK NPARTCRDLKMCHSDWKSGEYWIDPNQGCNLDAIKVYCNMETGQTCVFPTQPSVPQKNWYISP NPKEKKHVWFGESMTDGFPFEYGSEGSDPADVAIQLTFLRLMSTEASQNITYHCKNSVAYMDQQ TGNLKKALLLQGSNEIELRGEGNSRFTYSTLVDGCTSHTGTWGKTVIEYKTTKTSRLPIIDVAPLDIG APDQEFGLDIGPACFV

**Human COL3A1 protein sequence:** MMSFVQKGSWLLLALLHPTIILAQQEAVEGGCSHLGQSYADRDVWKPEPCQICVCDSGSVLCDDI ICDDQELDCPNPEIPFGECCAVCPQPPTAPTRPPNGQGPQGPKGDPGPPGIPGRNGDPGIPGQP GSPGSPGPPGICESCPTGPQNYSPQYDSYDVKSGVAVGGLAGYPGPAGPPGPPGPPGTSGHPG SPGSPGYQGPPGEPGQAGPSGPPGPPGAIGPSGPAGKDGESGRPGRPGERGLPGPPGIKGPA GIPGFPGMKGHRGFDGRNGEKGETGAPGLKGENGLPGENGAPGPMGPRGAPGERGRPGLPGA AGARGNDGARGSDGQPGPPGPPGTAGFPGSPGAKGEVGPAGSPGSNGAPGQRGEPGPQGHA GAQGPPGPPGINGSPGGKGEMGPAGIPGAPGLMGARGPPGPAGANGAPGLRGGAGEPGKNG AKGEPGPRGERGEAGIPGVPGAKGEDGKDGSPGEPGANGLPGAAGERGAPGFRGPAGPNGIPG EKGPAGERGAPGPAGPRGAAGEPGRDGVPGGPGMRGMPGSPGGPGSDGKPGPPGSQGESG RPGPPGPSGPRGQPGVMGFPGPKGNDGAPGKNGERGGPGGPGPQGPPGKNGETGPQGPPG PTGPGGDKGDTGPPGPQGLQGLPGTGGPPGENGKPGEPGPKGDAGAPGAPGGKGDAGAPGE RGPPGLAGAPGLRGGAGPPGPEGGKGAAGPPGPPGAAGTPGLQGMPGERGGLGSPGPKGDK GEPGGPGADGVPGKDGPRGPTGPIGPPGPAGQPGDKGEGGAPGLPGIAGPRGSPGERGETGP PGPAGFPGAPGQNGEPGGKGERGAPGEKGEGGPPGVAGPPGGSGPAGPPGPQGVKGERGSP GGPGAAGFPGARGLPGPPGSNGNPGPPGPSGSPGKDGPPGPAGNTGAPGSPGVSGPKGDAG QPGEKGSPGAQGPPGAPGPLGIAGITGARGLAGPPGMPGPRGSPGPQGVKGESGKPGANGLS GERGPPGPQGLPGLAGTAGEPGRDGNPGSDGLPGRDGSPGGKGDRGENGSPGAPGAPGHPG PPGPVGPAGKSGDRGESGPAGPAGAPGPAGSRGAPGPQGPRGDKGETGERGAAGIKGHRGFP GNPGAPGSPGPAGQQGAIGSPGPAGPRGPVGPSGPPGKDGTSGHPGPIGPPGPRGNRGERGS EGSPGHPGQPGPPGPPGAPGPCCGGVGAAAIAGIGGEKAGGFAPYYGDEPMDFKINTDEIMTSL KSVNGQIESLISPDGSRKNPARNCRDLKFCHPELKSGEYWVDPNQGCKLDAIKVFCNMETGETCI SANPLNVPRKHWWTDSSAEKKHVWFGESMDGGFQFSYGNPELPEDVLDVQLAFLRLLSSRASQ NITYHCKNSIAYMDQASGNVKKALKLMGSNEGEFKAEGNSKFTYTVLEDGCTKHTGEWSKTVFEY RTRKAVRLPIVDIAPYDIGGPDQEFGVDVGPVCFL

**Mouse COL3A1 protein sequence:** MMSFVQSGTWFLLTLLHPTLILAQQSNVDELGCSHLGQSYESRDVWKPEPCQICVCDSGSVLCD DIICDEEPLDCPNPEIPFGECCAICPQPSTPAPVLPDGHGPQGPKGDPGPPGIPGRNGDPGLPGQ PGLPGPPGSPGICESCPTGGQNYSPQFDSYDVKSGVGGMGGYPGPAGPPGPPGPPGSSGHPG SPGSPGYQGPPGEPGQAGPAGPPGPPGALGPAGPAGKDGESGRPGRPGERGLPGPPGIKGPA GMPGFPGMKGHRGFDGRNGEKGETGAPGLKGENGLPGDNGAPGPMGPRGAPGERGRPGLPG AAGARGNDGARGSDGQPGPPGPPGTAGFPGSPGAKGEVGPAGSPGSNGSPGQRGEPGPQGH AGAQGPPGPPGNNGSPGGKGEMGPAGIPGAPGLIGARGPPGPAGTNGIPGTRGPSGEPGKNGA KGEPGARGERGEAGSPGIPGPKGEDGKDGSPGEPGANGLPGAAGERGPSGFRGPAGPNGIPGE KGPPGERGGPGPAGPRGVAGEPGRDGTPGGPGIRGMPGSPGGPGNDGKPGPPGSQGESGRP GPPGPSGPRGQPGVMGFPGPKGNDGAPGKNGERGGPGGPGLPGPAGKNGETGPQGPPGPTG PAGDKGDSGPPGPQGLQGIPGTGGPPGENGKPGEPGPKGEVGAPGAPGGKGDSGAPGERGPP GTAGIPGARGGAGPPGPEGGKGPAGPPGPPGASGSPGLQGMPGERGGPGSPGPKGEKGEPG GAGADGVPGKDGPRGPAGPIGPPGPAGQPGDKGEGGSPGLPGIAGPRGGPGERGEHGPPGPA GFPGAPGQNGEPGAKGERGAPGEKGEGGPPGPAGPTGSSGPAGPPGPQGVKGERGSPGGPG TAGFPGGRGLPGPPGNNGNPGPPGPSGAPGKDGPPGPAGNSGSPGNPGIAGPKGDAGQPGEK GPPGAQGPPGSPGPLGIAGLTGARGLAGPPGMPGPRGSPGPQGIKGESGKPGASGHNGERGP PGPQGLPGQPGTAGEPGRDGNPGSDGQPGRDGSPGGKGDRGENGSPGAPGAPGHPGPPGPV GPSGKSGDRGETGPAGPSGAPGPAGARGAPGPQGPRGDKGETGERGSNGIKGHRGFPGNPGP PGSPGAAGHQGAIGSPGPAGPRGPVGPHGPPGKDGTSGHPGPIGPPGPRGNRGERGSEGSPG HPGQPGPPGPPGAPGPCCGGGAAAIAGVGGEKSGGFSPYYGDDPMDFKINTEEIMSSLKSVNG QIESLISPDGSRKNPARNCRDLKFCHPELKSGEYWVDPNQGCKMDAIKVFCNMETGETCINASP MTVPRKHWWTDSGAEKKHVWFGESMNGGFQFSYGPPDLPEDVVDVQLAFLRLLSSRASQNITY HCKNSIAYMDQASGNVKKSLKLMGSNEGEFKAEGNSKFTYTVLEDGCTKHTGEWSKTVFEYQTR KAMRLPIIDIAPYDIGGPDQEFGVDIGPVCFL

**Human LTBP2 protein sequence:** MRPRTKARSPGRALRNPWRGFLPLTLALFVGAGHAQRDPVGRYEPAGGDANRLRRPGGSYPAA AAAKVYSLFREQDAPVAGLQPVERAQPGWGSPRRPTEAEARRPSRAQQSRRVQPPAQTRRSTPL GQQQPAPRTRAAPALPRLGTPQRSGAAPPTPPRGRLTGRNVCGGQCCPGWTTANSTNHCIKPV CEPPCQNRGSCSRPQLCVCRSGFRGARCEEVIPDEEFDPQNSRLAPRRWAERSPNLRRSSAAG EGTLARAQPPAPQSPPAPQSPPAGTLSGLSQTHPSQQHVGLSRTVRLHPTATASSQLSSNALPP GPGLEQRDGTQQAVPLEHPSSPWGLNLTEKIKKIKIVFTPTICKQTCARGHCANSCERGDTTTLYS QGGHGHDPKSGFRIYFCQIPCLNGGRCIGRDECWCPANSTGKFCHLPIPQPDREPPGRGSRPRA LLEAPLKQSTFTLPLSNQLASVNPSLVKVHIHHPPEASVQIHQVAQVRGGVEEALVENSVETRPPP WLPASPGHSLWDSNNIPARSGEPPRPLPPAAPRPRGLLGRCYLNTVNGQCANPLLELTTQEDCC GSVGAFWGVTLCAPCPPRPASPVIENGQLECPQGYKRLNLTHCQDINECLTLGLCKDAECVNTR GSYLCTCRPGLMLDPSRSRCVSDKAISMLQGLCYRSLGPGTCTLPLAQRITKQICCCSRVGKAW GSECEKCPLPGTEAFREICPAGHGYTYASSDIRLSMRKAEEEELARPPREQGQRSSGALPGPAER QPLRVVTDTWLEAGTIPDKGDSQAGQVTTSVTHAPAWVTGNATTPPMPEQGIAEIQEEQVTPSTD VLVTLSTPGIDRCAAGATNVCGPGTCVNLPDGYRCVCSPGYQLHPSQAYCTDDNECLRDPCKGK GRCINRVGSYSCFCYPGYTLATSGATQECQDINECEQPGVCSGGQCTNTEGSYHCECDQGYIMV RKGHCQDINECRHPGTCPDGRCVNSPGSYTCLACEEGYRGQSGSCVDVNECLTPGVCAHGKCT NLEGSFRCSCEQGYEVTSDEKGCQDVDECASRASCPTGLCLNTEGSFACSACENGYWVNEDGT ACEDLDECAFPGVCPSGVCTNTAGSFSCKDCDGGYRPSPLGDSCEDVDECEDPQSSCLGGECK NTVGSYQCLCPQGFQLANGTVCEDVNECMGEEHCAPHGECLNSHGSFFCLCAPGFVSAEGGTS CQDVDECATTDPCVGGHCVNTEGSFNCLCETGFQPSPESGECVDIDECEDYGDPVCGTWKCEN SPGSYRCVLGCQPGFHMAPNGDCIDIDECANDTMCGSHGFCDNTDGSFRCLCDQGFEISPSGW DCVDVNECELMLAVCGAALCENVEGSFLCLCASDLEEYDAQEGHCRPRGAGGQSMSEAPTGDH APAPTRMDCYSGQKGHAPCSSVLGRNTTQAECCCTQGASWGDACDLCPSEDSAEFSEICPSGK GYIPVEGAWTFGQTMYTDADECVIFGPGLCPNGRCLNTVPGYVCLCNPGFHYDASHKKCEDHDE CQDLACENGECVNTEGSFHCFCSPPLTLDLSQQRCMNSTSSTEDLPDHDIHMDICWKKVTNDVC SEPLRGHRTTYTECCCQDGEAWSQQCALCPPRSSEVYAQLCNVARIEAEREAGVHFRPGYEYGP GPDDLHYSIYGPDGAPFYNYLGPEDTVPEPAFPNTAGHSADRTPILESPLQPSELQPHYVASHPEP PAGFEGLQAEECGILNGCENGRCVRVREGYTCDCFEGFQLDAAHMACVDVNECDDLNGPAVLCV HGYCENTEGSYRCHCSPGYVAEAGPPHCTAKE

**Mouse LTBP2 protein sequence:** MRAPTTARCSGCIRRVRWRGFLPLVLAVLMGTSHAQRDSIGRYEPASRDANRLWHPVGSHPAAA AAKVYSLFREPDAPVPGLSPSEWNQPAQGNPGRLAEAEARRPPRTQQLRRVQPPVQTRRSHPR GQQQIAARAAPSVARLETPQRPAAARRGRLTGRNVCGGQCCPGWTTSNSTNHCIKPVCQPPCQ NRGSCSRPQVCICRSGFRGARCEEVIPEEEFDPQNARPVPRRSVERAPGPHRSSEARGSLVTRIQ PLVPPPSPPPSRRLSQPWPLQQHSGPSRTVRRYPATGANGQLMSNALPSGLELRDSSPQAAHV NHLSPPWGLNLTEKIKKIKVVFTPTICKQTCARGRCANSCEKGDTTTLYSQGGHGHDPKSGFRIYF CQIPCLNGGRCIGRDECWCPANSTGKFCHLPVPQPDREPAGRGSRHRTLLEGPLKQSTFTLPLS NQLASVNPSLVKVQIHHPPEASVQIHQVARVRGELDPVLEDNSVETRASRRPHGNLGHSPWASNS IPARAGEAPRPPPVLSRHYGLLGQCYLSTVNGQCANPLGELTSQEDCCGSVGTFWGVTSCAPCP PRPAFPVIENGQLECPQGYKRLNLSHCQDINECLTLGLCKDSECVNTRGSYLCTCRPGLMLDPSR SRCVSDKAVSMQQGLCYRSLGSGTCTLPLVHRITKQICCCSRVGKAWGSTCEQCPLPGTEAFREI CPAGHGYTYSSSDIRLSMRKAEEEELASPLREQTEQSTAPPPGQAERQPLRAATATWIEAETLPDK GDSRAVQITTSAPHLPARVPGDATGRPAPSLPGQGIPESPAEEQVIPSSDVLVTHSPPDFDPCFAG ASNICGPGTCVSLPNGYRCVCSPGYQLHPSQDYCTDDNECMRNPCEGRGRCVNSVGSYSCLCY PGYTLVTLRDTQECQDIDECEQPGVCSGGRCSNTEGSYHCECDRGYIMVRKGHCQDINECRHPG TCPDGRCVNSPGSYTCLACEEGYVGQSGSCVDVNECLTPGICTHGRCINMEGSFRCSCEPGYEV TPDKKGCRDVDECASRASCPTGLCLNTEGSFTCSACQSGYWVNEDGTACEDLDECAFPGVCPT GVCTNTVGSFSCKDCDRGYRPNPLGNRCEDVDECEGPQSSCRGGECKNTEGSYQCLCHQGFQ LVNGTMCEDVNECVGEEHCAPHGECLNSLGSFFCLCAPGFASAEGGTRCQDVDECAATDPCPG GHCVNTEGSFSCLCETGFQPSPDSGECLDIDECEDREDPVCGAWRCENSPGSYRCILDCQPGFY VAPNGDCIDIDECANDTVCGNHGFCDNTDGSFRCLCDQGFETSPSGWECVDVNECELMMAVCG DALCENVEGSFLCLCASDLEEYDAEEGHCRPRVAGAQRIPEVRTEDQAPSLIRMECYSEHNGGPP CSQILGQNSTQAECCCTQGARWGKACAPCPSEDSVEFSQLCPSGQGYIPVEGAWTFGQTMYTD ADECVLFGPALCQNGRCLNIVPGYICLCNPGYHYDASSRKCQDHNECQDLACENGECVNTEGSF HCLCNPPLTLDLSGQRCVNSTSSTEDFPDHDIHMDICWKKVTNDVCSQPLRGHHTTYTECCCQD GEAWSQQCALCPPRSSEVYAQLCNVARIEAERGAGIHFRPGYEYGPGLDDLPENLYGPDGAPFY NYLGPEDTAPEPPFSNPASQPGDNTPVLEPPLQPSELQPHYLASHSEPLASFEGLQAEECGILNG CENGRCVRVREGYTCDCFEGFQLDAAHMACVDVNECEDLNGPAALCAHGHCENTEGSYRCHC SPGYVAEPGPPHCAAKE

**Human CKAP4 protein sequence:** MPSAKQRGSKGGHGAASPSEKGAHPSGGADDVAKKPPPAPQQPPPPPAPHPQQHPQQHPQN QAHGKGGHRGGGGGGGKSSSSSSASAAAAAAAASSSASCSRRLGRALNFLFYLALVAAAAFSG WCVHHVLEEVQQVRRSHQDFSRQREELGQGLQGVEQKVQSLQATFGTFESILRSSQHKQDLTEK AVKQGESEVSRISEVLQKLQNEILKDLSDGIHVVKDARERDFTSLENTVEERLTELTKSINDNIAIFTE VQKRSQKEINDMKAKVASLEESEGNKQDLKALKEAVKEIQTSAKSREWDMEALRSTLQTMESDIY TEVRELVSLKQEQQAFKEAADTERLALQALTEKLLRSEESVSRLPEEIRRLEEELRQLKSDSHGPKE DGGFRHSEAFEALQQKSQGLDSRLQHVEDGVLSMQVASARQTESLESLLSKSQEHEQRLAALQG RLEGLGSSEADQDGLASTVRSLGETQLVLYGDVEELKRSVGELPSTVESLQKVQEQVHTLLSQDQ AQAARLPPQDFLDRLSSLDNLKASVSQVEADLKMLRTAVDSLVAYSVKIETNENNLESAKGLLDDL RNDLDRLFVKVEKIHEKV

**Mouse CKAP4 protein sequence:** MPSAKQRGSKGGHGAASPSDKGAHPSGGADDVAKKPPAAPQQPQPPAPHPPQHPQNQAHRG GHRGRSSAATANASSASCSRRLGRVLNFLFYLSLVAAAAFSGWYVHHVLEEVQQVRRGHQDFSR QRDELGQGLQGVEQKVQSLQATFGTFESLLRNSQHKQDLTEKAVKEGESELNRISEVLQKLQNEIL KDLSDGIHVVKDARERDFTSLENTVEERLTELTKSINDNIAIFTDVQKRSQKEINEVKMKVASLEESK GDRSQDVKTLKDAVKEVQASMMSRERDIEALKSSLQTMESDVYTEVRELVSLKQEQQAFKQAAD SERLALQALTEKLLRSEESSSRLPEDIRRLEEELQQLKVGAHGSEEGAVFKDSKALEELQRQIEGLG ARLQYVEDGVYSMQVASARHTESLESLLSKSQEYEQRLAMLQEHVGNLGSSSDLASTVRSLGET QLALSSDLKELKQSLGELPGTVESLQEQVLSLLSQDQAQAEGLPPQDFLDRLSSLDNLKSSVSQV ESDLKMLRTAVDSLVAYSVKIETNENNLESAKGLLDDLRNDLDRLFLKVEKIHEKI

## REFERENCES

1. Davis J and Molkentin JD. Myofibroblasts: trust your heart and let fate decide. J Mol Cell Cardiol. 2014;70:9–18.

2. Travers JG, Kamal FA, Robbins J, Yutzey KE and Blaxall BC. Cardiac Fibrosis: The Fibroblast Awakens. Circ Res. 2016;118:1021–40.

3. Liu Y, Beyer A and Aebersold R. On the Dependency of Cellular Protein Levels on mRNA Abundance. Cell. 2016;165:535–50.

4. Schafer S, Viswanathan S, Widjaja AA, Lim WW, Moreno-Moral A, DeLaughter DM, Ng B, Patone G, Chow K, Khin E, Tan J, Chothani SP, Ye L, Rackham OJL, Ko NSJ, Sahib NE, Pua CJ, Zhen NTG, Xie C, Wang M, Maatz H, Lim S, Saar K, Blachut S, Petretto E, Schmidt S, Putoczki T, Guimaraes-Camboa N, Wakimoto H, van Heesch S, Sigmundsson K, Lim SL, Soon JL, Chao VTT, Chua YL, Tan TE, Evans SM, Loh YJ, Jamal MH, Ong KK, Chua KC, Ong BH, Chakaramakkil MJ, Seidman JG, Seidman CE, Hubner N, Sin KYK and Cook SA. IL-11 is a crucial determinant of cardiovascular fibrosis. Nature. 2017;552:110–115.

5. Chothani S, Schafer S, Adami E, Viswanathan S, Widjaja AA, Langley SR, Tan J, Wang M, Quaife NM, Pua CJ, D’Agostino G, Shekeran SG, George BL, Lim S, Cao EY, van Heesch S, Witte F, Felkin LE, Christodoulou EG, Dong J, Blachut S, Patone G, Barton PJR, Hubner N, Cook SA and Rackham OJL. Widespread Translational Control of Fibrosis in the Human Heart by RNA-Binding Proteins. Circulation. 2019.

6. van Heesch S, Witte F, Schneider-Lunitz V, Schulz JF, Adami E, Faber AB, Kirchner M, Maatz H, Blachut S, Sandmann CL, Kanda M, Worth CL, Schafer S, Calviello L, Merriott R, Patone G, Hummel O, Wyler E, Obermayer B, Mucke MB, Lindberg EL, Trnka F, Memczak S, Schilling M, Felkin LE, Barton PJR, Quaife NM, Vanezis K, Diecke S, Mukai M, Mah N, Oh SJ, Kurtz A, Schramm C, Schwinge D, Sebode M, Harakalova M, Asselbergs FW, Vink A, de Weger RA, Viswanathan S, Widjaja AA, Gartner-Rommel A, Milting H, Dos Remedios C, Knosalla C, Mertins P, Landthaler M, Vingron M, Linke WA, Seidman JG, Seidman CE, Rajewsky N, Ohler U, Cook SA and Hubner N. The Translational Landscape of the Human Heart. Cell. 2019;178:242–260 e29.

7. Kim S and Coulombe PA. Emerging role for the cytoskeleton as an organizer and regulator of translation. Nat Rev Mol Cell Biol. 2010;11:75–81.

8. Yao P and Fox PL. Aminoacyl-tRNA synthetases in medicine and disease. EMBO Mol Med. 2013;5:332–43.

9. Yao P, Poruri K, Martinis SA and Fox PL. Non-catalytic Regulation of Gene Expression by Aminoacyl-tRNA Synthetases. Top Curr Chem. 2013.

10. Arif A, Yao P, Terenzi F, Jia J, Ray PS and Fox PL. The GAIT translational control system. Wiley Interdiscip Rev RNA. 2018;9.

11. Mendes MI, Gutierrez Salazar M, Guerrero K, Thiffault I, Salomons GS, Gauquelin L, Tran LT, Forget D, Gauthier MS, Waisfisz Q, Smith DEC, Simons C, van der Knaap MS, Marquardt I, Lemes A, Mierzewska H, Weschke B, Koehler W, Coulombe B, Wolf NI and Bernard G. Bi-allelic Mutations in EPRS, Encoding the Glutamyl-Prolyl-Aminoacyl-tRNA Synthetase, Cause a Hypomyelinating Leukodystrophy. Am J Hum Genet. 2018;102:676–684.

12. Keller TL, Zocco D, Sundrud MS, Hendrick M, Edenius M, Yum J, Kim YJ, Lee HK, Cortese JF, Wirth DF, Dignam JD, Rao A, Yeo CY, Mazitschek R and Whitman M. Halofuginone and other febrifugine derivatives inhibit prolyl-tRNA synthetase. Nat Chem Biol. 2012;8:311–7.

13. Zhou H, Sun L, Yang XL and Schimmel P. ATP-directed capture of bioactive herbal-based medicine on human tRNA synthetase. Nature. 2013;494:121–4.

14. Kwon NH, Fox PL and Kim S. Aminoacyl-tRNA synthetases as therapeutic targets. Nature reviews Drug discovery. 2019.

15. Qin P, Arabacilar P, Bernard RE, Bao W, Olzinski AR, Guo Y, Lal H, Eisennagel SH, Platchek MC, Xie W, Del Rosario J, Nayal M, Lu Q, Roethke T, Schnackenberg CG, Wright F, Quaile MP, Halsey WS, Hughes AM, Sathe GM, Livi GP, Kirkpatrick RB, Qu XA, Rajpal DK, Faelth Savitski M, Bantscheff M, Joberty G, Bergamini G, Force TL, Gatto GJ, Jr., Hu E and Willette RN. Activation of the Amino Acid Response Pathway Blunts the Effects of Cardiac Stress. J Am Heart Assoc. 2017;6.

16. Sundrud MS, Koralov SB, Feuerer M, Calado DP, Kozhaya AE, Rhule-Smith A, Lefebvre RE, Unutmaz D, Mazitschek R, Waldner H, Whitman M, Keller T and Rao A. Halofuginone inhibits TH17 cell differentiation by activating the amino acid starvation response. Science. 2009;324:1334–8.

17. Khajuria RK, Munschauer M, Ulirsch JC, Fiorini C, Ludwig LS, McFarland SK, Abdulhay NJ, Specht H, Keshishian H, Mani DR, Jovanovic M, Ellis SR, Fulco CP, Engreitz JM, Schutz S, Lian J, Gripp KW, Weinberg OK, Pinkus GS, Gehrke L, Regev A, Lander ES, Gazda HT, Lee WY, Panse VG, Carr SA and Sankaran VG. Ribosome Levels Selectively Regulate Translation and Lineage Commitment in Human Hematopoiesis. Cell. 2018;173:90–103 e19.

18. Song DG, Kim D, Jung JW, Nam SH, Kim JE, Kim HJ, Kim JH, Lee SJ, Pan CH, Kim S and Lee JW. Glutamyl-prolyl-tRNA synthetase induces fibrotic extracellular matrix via both transcriptional and translational mechanisms. FASEB J. 2019;33:4341–4354.

19. Song DG, Kim D, Jung JW, Nam SH, Kim JE, Kim HJ, Kim JH, Pan CH, Kim S and Lee JW. Glutamyl-Prolyl-tRNA Synthetase Regulates Epithelial Expression of Mesenchymal Markers and Extracellular Matrix Proteins: Implications for Idiopathic Pulmonary Fibrosis. Front Pharmacol. 2018;9:1337.

20. Yao P, Wu J, Lindner D and Fox PL. Interplay between miR-574-3p and hnRNP L regulates VEGFA mRNA translation and tumorigenesis. Nucleic Acids Res. 2017;45:7950–7964.

21. Teekakirikul P, Eminaga S, Toka O, Alcalai R, Wang L, Wakimoto H, Nayor M, Konno T, Gorham JM, Wolf CM, Kim JB, Schmitt JP, Molkentin JD, Norris RA, Tager AM, Hoffman SR, Markwald RR, Seidman CE and Seidman JG. Cardiac fibrosis in mice with hypertrophic cardiomyopathy is mediated by non-myocyte proliferation and requires Tgf-beta. J Clin Invest. 2010;120:3520–9.

22. Da M, Feng Y, Xu J, Hu Y, Lin Y, Ni B, Qian B, Hu Z and Mo X. Association of aminoacyl-tRNA synthetases gene polymorphisms with the risk of congenital heart disease in the Chinese Han population. PLoS One. 2014;9:e110072.

23. Rau CD, Romay MC, Tuteryan M, Wang JJ, Santolini M, Ren S, Karma A, Weiss JN, Wang Y and Lusis AJ. Systems Genetics Approach Identifies Gene Pathways and Adamts2 as Drivers of Isoproterenol-Induced Cardiac Hypertrophy and Cardiomyopathy in Mice. Cell Syst. 2017;4:121–128 e4.

24. Galindo CL, Skinner MA, Errami M, Olson LD, Watson DA, Li J, McCormick JF, McIver LJ, Kumar NM, Pham TQ and Garner HR. Transcriptional profile of isoproterenol-induced cardiomyopathy and comparison to exercise-induced cardiac hypertrophy and human cardiac failure. BMC Physiol. 2009;9:23.

25. Robinson MM, Dasari S, Konopka AR, Johnson ML, Manjunatha S, Esponda RR, Carter RE, Lanza IR and Nair KS. Enhanced Protein Translation Underlies Improved Metabolic and Physical Adaptations to Different Exercise Training Modes in Young and Old Humans. Cell Metab. 2017;25:581–592.

26. Schuller AP, Wu CC, Dever TE, Buskirk AR and Green R. eIF5A Functions Globally in Translation Elongation and Termination. Mol Cell. 2017;66:194–205 e5.

27. Doerfel LK, Wohlgemuth I, Kothe C, Peske F, Urlaub H and Rodnina MV. EF-P is essential for rapid synthesis of proteins containing consecutive proline residues. Science. 2013;339:85–8.

28. Buskirk AR and Green R. Ribosome pausing, arrest and rescue in bacteria and eukaryotes. Philos Trans R Soc Lond B Biol Sci. 2017;372.

29. Mandal A, Mandal S and Park MH. Genome-wide analyses and functional classification of proline repeat-rich proteins: potential role of eIF5A in eukaryotic evolution. PLoS One. 2014;9:e111800.

30. UniProt C. UniProt: a worldwide hub of protein knowledge. Nucleic Acids Res. 2019;47:D506–D515.

31. Ishimura R, Nagy G, Dotu I, Zhou H, Yang XL, Schimmel P, Senju S, Nishimura Y, Chuang JH and Ackerman SL. RNA function. Ribosome stalling induced by mutation of a CNS-specific tRNA causes neurodegeneration. Science. 2014;345:455–9.

32. Park S, Ranjbarvaziri S, Lay FD, Zhao P, Miller MJ, Dhaliwal JS, Huertas-Vazquez A, Wu X, Qiao R, Soffer JM, Rau C, Wang Y, Mikkola HKA, Lusis AJ and Ardehali R. Genetic Regulation of Fibroblast Activation and Proliferation in Cardiac Fibrosis. Circulation. 2018;138:1224–1235.

33. Sala-Newby GB, George SJ, Bond M, Dhoot GK and Newby AC. Regulation of vascular smooth muscle cell proliferation, migration and death by heparan sulfate 6-O-endosulfatase1. FEBS Lett. 2005;579:6493–8.

34. Gorsi B, Liu F, Ma X, Chico TJ, v A, Kramer KL, Bridges E, Monteiro R, Harris AL, Patient R and Stringer SE. The heparan sulfate editing enzyme Sulf1 plays a novel role in zebrafish VegfA mediated arterial venous identity. Angiogenesis. 2014;17:77–91.

35. Liu Y, Morley M, Brandimarto J, Hannenhalli S, Hu Y, Ashley EA, Tang WH, Moravec CS, Margulies KB, Cappola TP, Li M and consortium MA. RNA-Seq identifies novel myocardial gene expression signatures of heart failure. Genomics. 2015;105:83–9.

36. Gladka MM, Molenaar B, de Ruiter H, van der Elst S, Tsui H, Versteeg D, Lacraz GPA, Huibers MMH, van Oudenaarden A and van Rooij E. Single-Cell Sequencing of the Healthy and Diseased Heart Reveals Cytoskeleton-Associated Protein 4 as a New Modulator of Fibroblasts Activation. Circulation. 2018;138:166–180.

37. Liu Z, Wang L, Welch JD, Ma H, Zhou Y, Vaseghi HR, Yu S, Wall JB, Alimohamadi S, Zheng M, Yin C, Shen W, Prins JF, Liu J and Qian L. Single-cell transcriptomics reconstructs fate conversion from fibroblast to cardiomyocyte. Nature. 2017;551:100–104.

38. Rockey DC, Bell PD and Hill JA. Fibrosis--a common pathway to organ injury and failure. N Engl J Med. 2015;372:1138–49.

39. van Riggelen J, Yetil A and Felsher DW. MYC as a regulator of ribosome biogenesis and protein synthesis. Nat Rev Cancer. 2010;10:301–9.

40. Qi F, Motz M, Jung K, Lassak J and Frishman D. Evolutionary analysis of polyproline motifs in Escherichia coli reveals their regulatory role in translation. PLoS Comput Biol. 2018;14:e1005987.

41. Guidotti G, Brambilla L and Rossi D. Cell-Penetrating Peptides: From Basic Research to Clinics. Trends Pharmacol Sci. 2017;38:406–424.

42. Yao P, Potdar AA, Arif A, Ray PS, Mukhopadhyay R, Willard B, Xu Y, Yan J, Saidel GM and Fox PL. Coding Region Polyadenylation Generates a Truncated tRNA Synthetase that Counters Translation Repression. Cell. 2012;149:88–100.

43. Ray PS, Jia J, Yao P, Majumder M, Hatzoglou M and Fox PL. A stress-responsive RNA switch regulates VEGFA expression. Nature. 2009;457:915–9.

44. Yao P, Potdar AA, Ray PS, Eswarappa SM, Flagg AC, Willard B and Fox PL. The HILDA complex coordinates a conditional switch in the 3’-untranslated region of the VEGFA mRNA. PLoS Biol. 2013;11:e1001635.

45. Venkata Subbaiah KC, Wu J, Potdar A and Yao P. hnRNP L-mediated RNA switches function as a hypoxia-induced translational regulon. Biochem Biophys Res Commun. 2019.

46. Megan E. Forrest AN, Thomas J. Sweet, Daniel Arango, Gavin Hanson, James Ellis, Shalini Oberdoerffer, Jeff Coller, Olivia S. Rissland. Codon usage and amino acid identity are major determinants of mRNA stability in humans. BioRxiv. 2018.

47. Wu Q, Medina SG, Kushawah G, DeVore ML, Castellano LA, Hand JM, Wright M and Bazzini AA. Translation affects mRNA stability in a codon-dependent manner in human cells. Elife. 2019;8.

48. Holst CR, Bou-Reslan H, Gore BB, Wong K, Grant D, Chalasani S, Carano RA, Frantz GD, Tessier-Lavigne M, Bolon B, French DM and Ashkenazi A. Secreted sulfatases Sulf1 and Sulf2 have overlapping yet essential roles in mouse neonatal survival. PLoS One. 2007;2:e575.

49. Ai X, Kitazawa T, Do AT, Kusche-Gullberg M, Labosky PA and Emerson CP, Jr. SULF1 and SULF2 regulate heparan sulfate-mediated GDNF signaling for esophageal innervation. Development. 2007;134:3327–38.

50. Ai X, Do AT, Kusche-Gullberg M, Lindahl U, Lu K and Emerson CP, Jr. Substrate specificity and domain functions of extracellular heparan sulfate 6-O-endosulfatases, QSulf1 and QSulf2. J Biol Chem. 2006;281:4969–76.

51. Yue X, Li X, Nguyen HT, Chin DR, Sullivan DE and Lasky JA. Transforming growth factor-beta1 induces heparan sulfate 6-O-endosulfatase 1 expression in vitro and in vivo. J Biol Chem. 2008;283:20397–407.

52. Korf-Klingebiel M, Reboll MR, Grote K, Schleiner H, Wang Y, Wu X, Klede S, Mikhed Y, Bauersachs J, Klintschar M, Rudat C, Kispert A, Niessen HW, Lubke T, Dierks T and Wollert KC. Heparan Sulfate-Editing Extracellular Sulfatases Enhance Vascular Endothelial Growth Factor Bioavailability for Ischemic Heart Repair. Circ Res. 2019.

## References

1. Skarnes WC, Rosen B, West AP, Koutsourakis M, Bushell W, Iyer V, Mujica AO, Thomas M, Harrow J, Cox T, Jackson D, Severin J, Biggs P, Fu J, Nefedov M, de Jong PJ, Stewart AF and Bradley A. A conditional knockout resource for the genome-wide study of mouse gene function. Nature. 2011;474:337–42.

2. Kanisicak O, Khalil H, Ivey MJ, Karch J, Maliken BD, Correll RN, Brody MJ, SC JL, Aronow BJ, Tallquist MD and Molkentin JD. Genetic lineage tracing defines myofibroblast origin and function in the injured heart. Nat Commun. 2016;7:12260.

3. Knight WE, Chen S, Zhang Y, Oikawa M, Wu M, Zhou Q, Miller CL, Cai Y, Mickelsen DM, Moravec C, Small EM, Abe J and Yan C. PDE1C deficiency antagonizes pathological cardiac remodeling and dysfunction. Proc Natl Acad Sci U S A. 2016.

4. Trembley MA, Quijada P, Agullo-Pascual E, Tylock KM, Colpan M, Dirkx RA, Jr., Myers JR, Mickelsen DM, de Mesy Bentley K, Rothenberg E, Moravec CS, Alexis JD, Gregorio CC, Dirksen RT, Delmar M and Small EM. Mechanosensitive Gene Regulation by Myocardin-Related Transcription Factors Is Required for Cardiomyocyte Integrity in Load-Induced Ventricular Hypertrophy. Circulation. 2018;138:1864–1878.

5. Yao P, Wu J, Lindner D and Fox PL. Interplay between miR-574-3p and hnRNP L regulates VEGFA mRNA translation and tumorigenesis. Nucleic Acids Res. 2017;45:7950–7964.

6. Bolger AM, Lohse M and Usadel B. Trimmomatic: a flexible trimmer for Illumina sequence data. Bioinformatics. 2014;30:2114–20.

7. Dobin A, Davis CA, Schlesinger F, Drenkow J, Zaleski C, Jha S, Batut P, Chaisson M and Gingeras TR. STAR: ultrafast universal RNA-seq aligner. Bioinformatics. 2013;29:15–21.

8. Liao Y, Smyth GK and Shi W. featureCounts: an efficient general purpose program for assigning sequence reads to genomic features. Bioinformatics. 2014;30:923–30.

9. Love MI, Huber W and Anders S. Moderated estimation of fold change and dispersion for RNA-seq data with DESeq2. Genome Biol. 2014;15:550.

10. Team RC. R: A language and environment for statistical computing. R Foundation for Statistical Computing, Vienna, Austria. URL https://www.R-project.org/. 2016.

11. Huang da W, Sherman BT and Lempicki RA. Systematic and integrative analysis of large gene lists using DAVID bioinformatics resources. Nat Protoc. 2009;4:44–57.

12. Mandal A, Mandal S and Park MH. Genome-wide analyses and functional classification of proline repeat-rich proteins: potential role of eIF5A in eukaryotic evolution. PLoS One. 2014;9:e111800.

